# Role of SARS-CoV-2 mutations in the evolution of the COVID-19 pandemic

**DOI:** 10.1101/2023.05.01.538506

**Authors:** Philippe Colson, Hervé Chaudet, Jérémy Delerce, Pierre Pontarotti, Anthony Levasseur, Jacques Fantini, Bernard LA Scola, Christian Devaux, Didier Raoult

## Abstract

RNA viruses, including SARS-CoV-2, evolve by mutation acquisition, or by hybridization between viral genomes. The SARS-CoV-2 pandemic provided an exceptional opportunity to analyze the mutations that appeared over a three-year period.

In this study, we analysed the type of mutations and their epidemic consequences on the thousands of genomes produced in our laboratory. These were obtained by next-generation sequencing from respiratory samples performed for genomic surveillance. The frequencies of mutations were calculated using Nextclade, Microsoft Excel, and an in-house Python script. In total, 61,397 genomes matching 483 Pangolin lineages were analyzed; 22,225 nucleotide mutations were identified, and of them 220 (1.0%) were each at the root of at least 836 genomes, a frequency threshold classifying mutations as “hyperfertile”. Two of these seeded the pandemic in Europe, namely a mutation in the RNA-dependent RNA polymerase associated with an increased mutation rate (P323L) and one in the spike protein (D614G), which plays a particular role in virus fitness. Most of these 220 “hyperfertile” mutations occurred in areas not predicted to be associated with increased virulence. Their number was 8±6 (0-22) per 1,000 nucleotides on average per gene. They were 3.7 times more frequent in accessory than informational genes (14 versus 4; p= 0.0037). Particularly, they were 4.1 times more frequent in ORF8 than in the gene encoding RNA polymerase. Interestingly, stop codons were present in 97 positions, almost only in six accessory genes including ORF7a (25 per 100 codons) and ORF8 (21). Furthermore, 1,661 mutations (16.3%) were associated with a lower number of “offspring” (50-835) and classified as “fertile”.

In conclusion, except for two initial mutations that could predict a change in the dynamics of the epidemic (mutation rate and change in the virus attachment site), most of the “hyperfertile” mutations did not predict the emergence of a new epidemic form. Significantly, some mutations were in non-coding areas and some consisted of stop codons, indicating that some genes (particularly ORF7a and ORF8) were rather “non-virulence genes” at a given stage of the epidemic, which is an unusual concept for viruses.

## INTRODUCTION

SARS-CoV-2 genomes are among the largest RNA virus genomes with approximately 30,000 nucleotides (Hartenian *et al*., 2020). They are, for instance, three times larger than the human immunodeficiency virus genome. At the time of emergence of SARS-CoV-2 it was considered that the mutation rate of coronaviruses must be relatively low in order to maintain the integrity of such a large genome (Worobey *et al*., 2020). It has been suggested that mutations were relatively rare, as these viruses benefit from an error correction system through a 3’-5’exonuclease (encoded by the NSP14 gene) (Rausch *et al*., 2020; Dearlove *et al*., 2020). Based on such a concept of probable genomic stability, pharmaceutic companies thought they could develop anti-spike protein vaccines aimed at controlling the spread of SARS-CoV-2 (Yang *et al*., 2020). But this hypothesis did not prevent the epidemiological molecular dynamic of the emergence and replacements of SARS-CoV-2 lineages (Carabelli *et al*., 2023). The succession of variants over time and the at least partial immune escape of the Omicron variants highlighted the need to gain a better understanding of the genomic evolution of SARS-CoV-2.

At our medical site dedicated to infectious diseases (the Méditerranée Infection university hospital institute (IHU)), we performed diagnosis, virus isolation, and next-generation sequencing (NGS) of SARS-CoV-2 genomes (La Scola *et al*., 2020; Colson *et al*., 2022). Thanks to the power of NGS, since the very beginning of the pandemic in our geographical area we were able to sequence more than 70,000 genomes (available in sequence databases). This allowed us to decipher the genetic evolution of SARS-CoV-2. The rapid evolution of this virus does not seem to be particularly affected by the human measures taken to control viral spread (Carabelli *et al*., 2023), and the apparent regularity of the mutations over time (https://nextstrain.org/; Markov *et al*., 2023) raises a fundamental question about the importance of the type of evolution proclaimed by the Red Queen theory (Van Valen, 1973), the Court Jester theory (Benton, 2009), or even the Mistigri theory, which we recently reported (Colson *et al*., 2023) (Box 1).

#### Box 1. Comparison of evolutionary models key characteristics

**The Red Queen hypothesis (**Van Valen, 1973)

Species must constantly proliferate and are strongly influenced by interactions among species (arms race for evolution). Extinction of a virus species occurs randomly with respect to time (any homogeneous group of organisms deteriorates at a stochastically constant rate), but non-randomly with respect to ecology (interspecific competition increases fitness and better fitness of one virus species deteriorates the fitness of coexisting virus species). Because coexisting species have a tendency to evolve in order to survive, no species gains a long-term increase in fitness. This intrinsic factor hypothesis was also illustrated by the “takes all the running you can do to keep in the same place” rule from the “Alice’s Adventures in Wonderland” book of Lewis Carroll (Carroll, 1865).

**The Court Jester hypothesis** (Barnosky, 2001; Benton, 2009)

Evolution, speciation and extinction rarely happen, except in response to unpredictable (stochastic) perturbations of the environment by abiotic factors changing the ground rules for the species and inducing a co-evolutionary switch in host-virus interactions. The time scale is likely longer than with the Red Queen model. This extrinsic factor hypothesis was also reported to be illustrated by the behaviour of the licensed fool of Medieval times.

**The Mistigri hypothesis (**Georgiades *et al*., 2011; Colson *et al*., 2023)

In a new ecological niche, the disappearance of genes (and associated functions) non-essential for the replication of the virus may be associated with improved fitness and/or virulence (Georgiades *et al*., 2011). This relates to the “use it or lose it” theory (Moran, 2002). This hypothesis was also reported to be illustrated by the “backpack rule”, which means: remove what you do not need to carry in your backpack before leaving and you can walk longer (Colson *et al*., 2023). As a matter of fact, the Mistigri rule relates to the game of cards for which the winners are those who get rid of the Mistigri (usually the jack of club) card (Colson et al., 2023).

Mutations within SARS-CoV-2 genomes were found to occur about once every 15 days (Carabelli *et al*., 2023) based on the analysis of more than 15 million viral genomes available worldwide as well as on our own genome database (Markov *et al*., 2023; Colson *et al*., 2022). In practice, these mutations can be of several kinds. Some are beneficial, generating a clone with at least 50 descendants (Colson *et al*., 2023), while others are of the neutral or weakly deleterious type, and their accumulation leads to a widening of the diversity of the initial clone and to an extinction of this clone (Elena *et al*., 2000; Colson *et al*., 2023). Finally, mutations can also be lethal, but those are not observable since the virus does not replicate. This phenomenon has been observed *in vitro* including with the vesicular stomatitis virus (Elena and Moya, 1999). By studying the Pangolin lineage B.1.160 (Marseille-4 variant) which predominated in France between August 2020 and February 2021, we were able to measure that, on average for a given variant, if there was no new beneficial mutation generating a new sub-variant, the accumulation of seven to eight non-beneficial mutations was associated with a funnel-like dispersion of the viral genomes, which was in turn associated with the decrease or even the disappearance of the viral clone (Colson *et al*., 2023). Here, we investigated a large set of 61,397 SARS-CoV-2 genomes sequenced in our institute covering the entire period of the pandemic from its onset, to get a better understanding of the frequency, nature and distribution of the mutations, and thereby of the evolution of SARS-CoV-2.

## MATERIALS AND METHODS

### Obtaining of SARS-CoV-2 genomes

All SARS-CoV-2 genomes analyzed in the present study were obtained by NGS at the Méditerranée Infection IHU, part of the university hospitals of Marseille (Assistance Publique-Hôpitaux de Marseille (AP-HM), Marseille, south-eastern France) from respiratory samples collected from patients between 24 February 2020 and 15 October 2022 for the purpose of diagnosing SARS-CoV-2 infection and sent to our clinical diagnosis laboratory. They were obtained in the framework of genomic surveillance, as recommended since 2021 by the French government (https://www.santepubliquefrance.fr/dossiers/coronavirus-covid-19/consortium-emergen), using the Illumina or Nanopore technologies, as previously reported (Colson *et al*., 2022; Colson *et al*., 2023). SARS-CoV-2 genomes were collected from the bioinformatics server of our institute where they had been stored.

### SARS-CoV-2 genome analysis

SARS-CoV-2 genomes were first reanalyzed using the Nextclade tool (https://clades.nextstrain.org/; Aksamentov *et al*., 2021). The resulting file was then manually curated using Microsoft Excel software (https://www.microsoft.com/fr-fr/microsoft-365/excel). Thus, genomes covering at least 90% of the size of the reference genome of the Wuhan-Hu-1 isolate with GenBank accession number NC_045512.2, and not obtained from a same patient and two samples fewer than 60 days apart (minimum period of time chosen by the eCDC to consider a viral re-infection (European Centre for Disease Prevention and Control, 2021) were selected. SARS-CoV-2 genomes conserved in the database analyzed in this study are available in the NCBI GenBank nucleotide sequence database (Sayers *et al*., 2023), in the GISAID database (https://gisaid.org/; Elbe *et al*., 2017), or on the IHU Méditerranée Infection website (https://www.mediterranee-infection.com/acces-ressources/donnees-pour-articles/) (see the Data Availability Statement).

### Frequencies of nucleotide and amino acid mutations

The total frequencies of nucleotide and amino acid mutations and the frequencies of these mutations according to Nextstrain clades (https://nextstrain.org/; Hadfield *et al*., 2018) and Pangolin lineages (https://cov-lineages.org/resources/pangolin.html; Rambaut *et al*., 2020) were calculated from the Nextclade output file (https://clades.nextstrain.org/; Aksamentov *et al*., 2021) generated by analyzing the whole set of SARS-CoV-2 genomes, using an in-house script written in Python (https://www.python.org/).

### Beneficial and neutral or deleterious mutations

We classified the mutations according to their frequencies among genomes. We considered that the most frequent mutations were beneficial for the viruses, whereas those with less frequency were deleterious or, at best, neutral. Furthermore, we determined a threshold of frequencies of mutations to delineate those which were highly frequent by searching for the most appropriate breakpoint in frequency distribution. We used the search algorithm for multiple structural changes in linear models in a two-fold procedure: a first search for the curve elbow, then a search for multiple changes within the half-curve corresponding to the most frequent mutations, with an appearance ordering of the points (Bai and Perron, 2003). We thus defined “hyperfertile” mutations as being the most frequent (so-named because they were associated with tremendous numbers of offspring) and “fertile” mutations as those of medium frequency (so-named because they were associated with large numbers of offspring). Neutral/deleterious mutations were defined as those of very low frequency, with an arbitrary threshold of 50 genomes harboring them.

### Definition of gene categories

SARS-CoV-2 gene categories were defined based on functions attributed to the proteins they encode. A total of 27 proteins have been annotated for SARS-CoV-2 (https://www.ncbi.nlm.nih.gov/data-hub/taxonomy/2697049/; Lubin *et al*., 2020; Jungreis *et al*., 2021; Prates *et al*., 2021) (Figure 1). The four categories of proteins were: (i) structural proteins; (ii) informational proteins (proteins involved in information storage and processing, which includes Clusters of Orthologous Groups (COGs) (https://www.ncbi.nlm.nih.gov/research/cog/) (Tatusov *et al*., 1997) of proteins with functional categories related to information storage and processing (J, A, K, L and B) and nucleotide transport and metabolism (F) (Boyer *et al*., 2010); (iii) other non-structural proteins; and (iv) so-called accessory proteins (Figure 1). Classification of the SARS-CoV-2 proteins was attempted by performing a similarity search with SARS-CoV-2 proteins recovered from GenBank (https://www.ncbi.nlm.nih.gov/data-hub/taxonomy/2697049/) in the COG database using the COG Guess web application (http://www-archbac.u-psud.fr/genomics/cog_guess.html).

**Figure 1.**
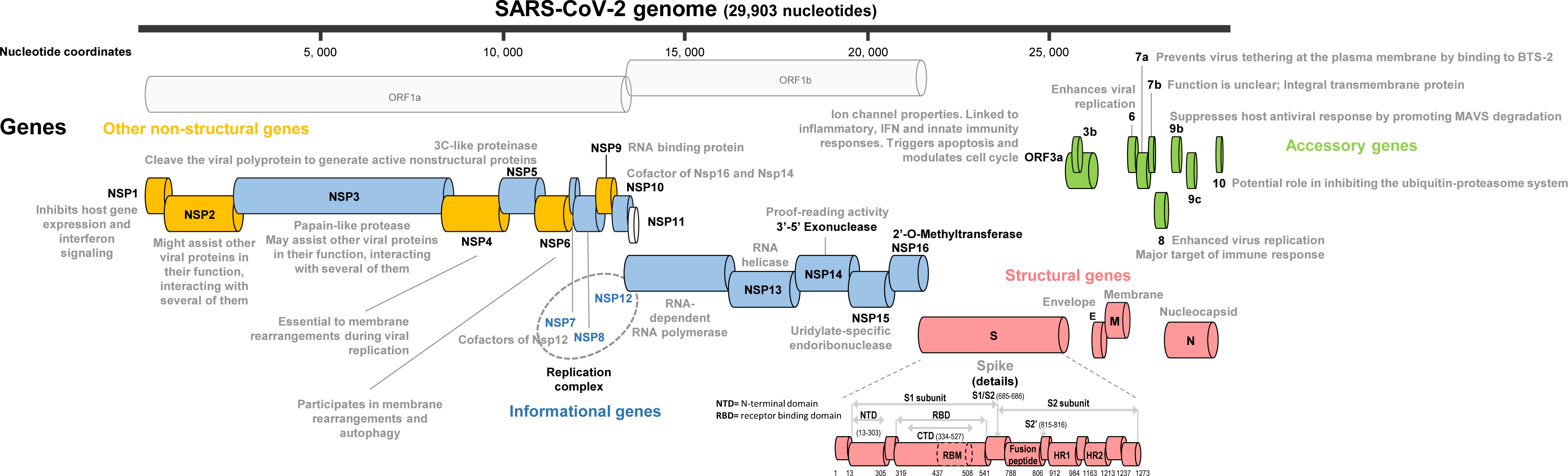
Map of the SARS-CoV-2 genome and genes. Functions of proteins were taken from Lubin *et al*., 2020; Jungreis *et al*., 2021; and Prates *et al*., 2021.

### Mutations per viral genes and gene categories

Nucleotide and amino acid mutations were counted for genes and gene categories. Gene positions were those available from the UCSC genome browser web application (https://genome.ucsc.edu/cgi-bin/hgGateway; Fernandes *et al*., 2020) for genome no. NC_045512.2. Correspondence between nucleotide and amino acid positions within the genome and genes was performed using Microsoft Excel. For comparison of frequencies of mutations, ORF3c, ORF3d, ORF9b, ORF9c, and ORF10 genes were not considered as they encode proteins considered as putative and/or shorter than 50 amino acids, or as they are an alternative reading frame of another gene.

### Protein structural analyses

Structural models of the spike protein and of the RNA-dependent RNA polymerase were generated from pdb files as previously reported (Colson *et al*., 2023). In the case of the spike protein, the gaps in the crystal structure were fixed by incorporating the missing amino acids with the Robetta protein structure prediction tool (Kim *et al*., 2004) before energy minimization with the Polak–Ribière algorithm of HyperChem was performed (Guex and Peitsch, 1997; Froimowitz, 1993). The structure of the P323L mutant of RNA-dependent RNA polymerase (amino acid residues 84-929) was derived from pdb file 7bv2 (Yin *et al*., 2020) with the same strategy. Mutant proteins were generated with Swiss-PdbViewer (Guex and Peitsch, 1997), and several rounds of energy minimization were carried out as previously reported (Di Scala and Fantini, 2017).

### Graphical representations and statistical analyses

Graphs were plotted using Microsoft Excel. Statistical analyses were performed using R software version 4.0.2 (https://cran.r-project.org/) and the Openepi online tool (https://www.openepi.com/). A *p* value <0.05 was considered as statistically significant.

## RESULTS

### Definition of gene categories

The 27 SARS-CoV-2 genes were classified as follows into four main categories: (i) four, namely S (spike), E (Envelope), M (Membrane), and N (Nucleocapsid), were “structural” genes; (ii) nine, namely NSP12 (RNA-dependent RNA polymerase), NSP13 (RNA helicase), NSP14 (3’-5’-exonuclease), NSP15 (Endoribouclease), NSP16 (2’-O-Methyltransferase), NSP7 and NSP8 (which are both involved in the replication complex with NSP12), NSP10 (cofactor of NSP14 and NSP16), and NSP3 and NSP5 (papain-like and 3C-like proteinases, respectively, that perform proteolytic cleavage of the polyproteins), were classified as “informational” genes; (iii) five, namely NSP1, NSP2, NSP4, NSP6, and NSP9, were “other non-structural” genes, which encode for proteins with various functions not precisely identified in some cases; and (iv) seven, namely ORF3a, ORF6, ORF7a, ORF7b, ORF8, ORF9b and ORF10, were “accessory” genes, which encode for proteins the functions of which were reported to modulate viral replication and interfere with host immune responses (https://www.ncbi.nlm.nih.gov/data-hub/taxonomy/2697049/; Lubin *et al*., 2020; Jungreis *et al*., 2021; Prates *et al*., 2021) (Figure 1).

### Number of SARS-CoV-2 genomes analyzed

A total of 61,397 genomic sequences were selected for the composition of the final curated set and were further analyzed. They had been obtained from patients sampled between 24 February 2020 and 17 October 2022 (a period of 32 months). For these genomes, the number of Nextclade lineages was 28, and the number of Pangolin lineages was 483 (Supplementary Table S1). The number of genomes per Nextclade lineage ranged from 1 to 20,168, and the number of genomes per Pangolin lineage ranged from 1 to 8,992. The time distribution of the main SARS-CoV-2 lineages based on the set of genomes analysed here and over the study period is shown in Supplementary Figure S1.

### Frequencies of mutations in SARS-CoV-2 genomes

Regarding nucleotide mutations, 22,225 different mutations were identified in ≥1 of the 61,397 SARS-CoV-2 genomes (Table 1; Supplementary Table S2; Supplementary Figure S2). A total of 21,525 mutations were substitutions, while 580 were deletions and 120 were insertions. These mutations stood at 16,692 nucleotide positions. A total of 12,015 positions harbored a single type of mutation while 4,677 positions harbored different types of mutations. A statistical analysis consisting in searching for breakpoints in the distribution of mutation frequencies enabled us to define that 220 mutations were “hyperfertile”, being harbored by ≥836 genomes (Figure 2; Supplementary Figure S3). In addition, 1,625 other mutations harbored by between 50 and 835 genomes were defined as “fertile”. Thus, “hyperfertile” and “fertile” nucleotide mutations accounted for 1.0% and 6.3% of the whole set of mutations, respectively, and stood at 1.3% and 8.2% of the mutated nucleotide positions. Four mutations were found in ≥50,000 genomes. They included three mutations that were present in SARS-CoV-2 that caused the first infections diagnosed in our geographical area as well as in the rest of France and Europe (Pachetti *et al*., 2020a; 2020b). These mutations are located in the genes encoding the spike (S gene; A23403G), generating D614G amino acid substitution (Supplementary Fig. S4), the RNA-dependent RNA polymerase (NSP12; C14408U), generating P323L amino acid substitution (Supplementary Fig. S4), and a papain-like proteinase with phosphoesterase activity (NSP3; C3037U). It is also worthy to note that among the mutations the most frequent, those in fourteenth and nineteenth positions by decreasing order of frequency were two mutations, A28271- or A28271U, which were previously reported as being located in a Kozak site of the N gene and as associated with the spread of Alpha, Delta and Omicron variants in combination with amino acid mutations P681H/R in the spike and R203K/M in the nucleocapsid (Yang *et al*., 2020).

**Figure 2.**
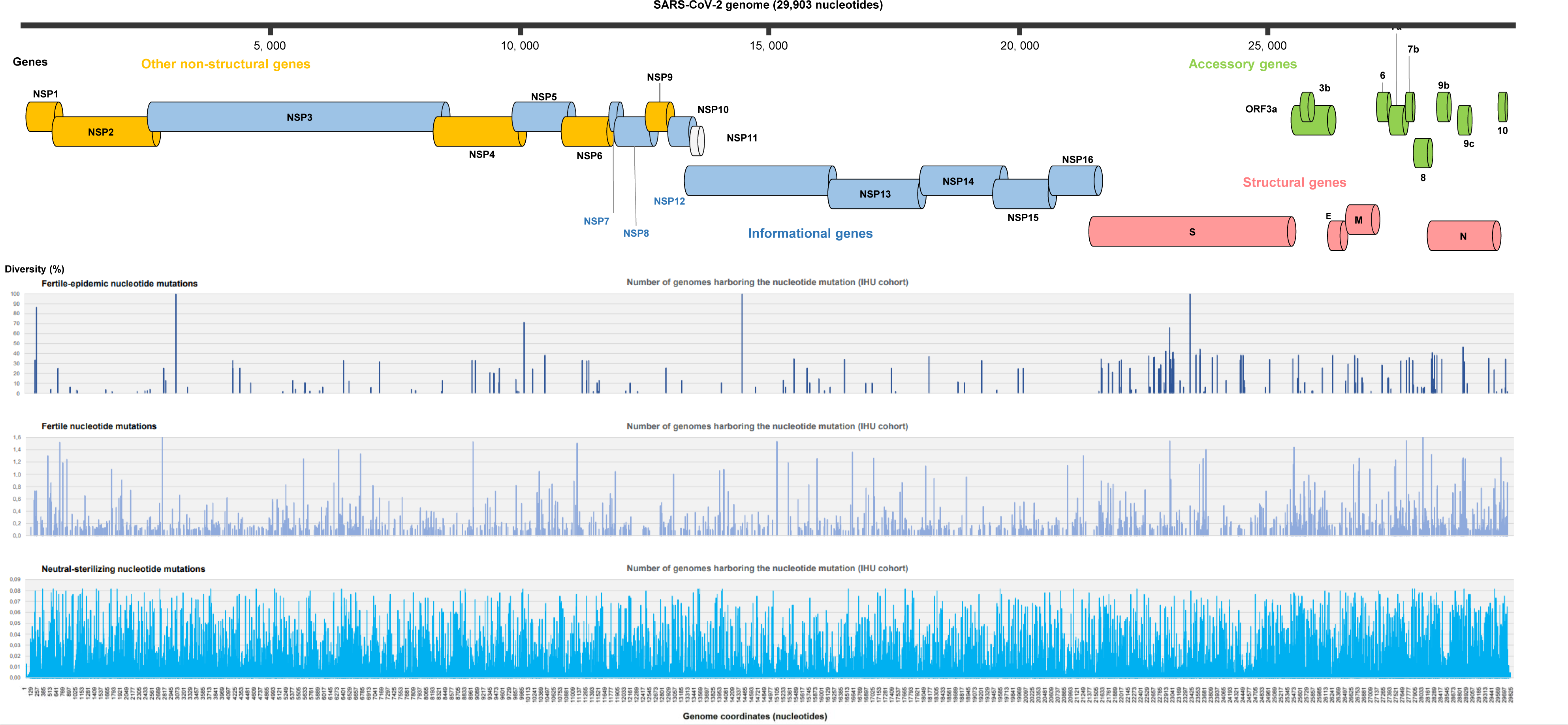
Distribution along the 61,397 SARS-CoV-2 genomes obtained in our institute of “hyperfertile”, “fertile”, and neutral or deleterious nucleotide mutations. IHU, University hospital institute Méditerranée Infection A map of the SARS-CoV-2 genome and genes is shown at the top of the figure.

**Table 1.** Classification of nucleotide and amino acid mutations in SARS-CoV-2 genomes from our institute based on their frequencies.

| Nature of mutations | Frequencies |  |  |  | Proportions (%) |  |  |
| --- | --- | --- | --- | --- | --- | --- | --- |
|  | Total | “Hyperfertile” | “Fertile” | Neutral or deleterious | “Hyperfertile” | “Fertile” | Neutral or deleterious |
| **Nucleotides** |  |  |  |  |  |  |  |
| Mutations | **22,221** | 220 | 1,400 | 20,601 | *1.0* | *6.3* | *92.7* |
| Mutated genome positions | **16,692** | 213 | 1,361 | 15,118 | *1.3* | *8.2* | *90.6* |
| **Amino acids** |  |  |  |  |  |  |  |
| Mutations | **13,224** | 191 | 755 | 12,278 | *1.4* | *5.7* | *92.9* |
| Mutated codons | **6,706** | 186 | 668 | 5,852 | *2.8* | *10.0* | *87.3* |
| **Stop codons** |  |  |  |  |  |  |  |
| According to classification of codon changes | **97** | 3 | 7 | 87 | *3.1* | *7.2* | *89.7* |
| According to distribution of stop codon frequencies | **97** | 5 | 31 | 61 | *5.2* | *32.0* | *62.9* |

Regarding amino acid mutations, 13,224 different mutations were identified in at least one gene product from at least one of the 61,397 SARS-CoV-2 genomes (Table 1; Supplementary Table S3). A total of 12,579 mutations were substitutions, while 597 were deletions and 48 were insertions. These mutations stood at 6,706 codons, and 2,901 of these codons included a single type of mutation, while several different mutations were identified at the 3,805 remaining mutated codons. A search for breakpoints in the distribution of mutation frequencies enabled us to define that 191 mutations were “hyperfertile”, being harbored by gene products from ≥628 genomes (Supplementary Figure S3). In addition, 755 other mutations harbored by gene products from between 50 and 627 genomes were defined as “fertile”. Thus, “hyperfertile” and “fertile” codon changes accounted for 1.4% and 5.7% of the whole set of codon changes, respectively, and stood at 2.8% and 10.0% of the mutated codons.

### Frequencies of mutations according to SARS-CoV-2 lineages

Considering the 483 Pangolin lineages, the mean proportion of these lineages for which at least one genome harbors a given mutation was 0.9% for nucleotide mutations and 1.0% for amino acid mutations, and maximum proportions were 98.0% both for nucleotide and amino acid mutations. In addition, the mean proportion of these lineages for which ≥50 genomes harbor a given mutation was 0.04% for nucleotide mutations and 1.0% for amino acid mutations, and the maximum proportions were 17.0% both for nucleotide and amino acid mutations. The genomes of six Pangolin lineages (namely BA.2 (Omicron 21L), AY.43 (Delta 21J), B.1.160 (Marseille-4), B.1.1.7 (Alpha), BA.1.1 (Omicron 21K) and BA.5.1 (Omicron 22B)) harbored >10% of all 22,225 nucleotide mutations observed in the whole set of genomes. Moreover, the genomes from five Pangolin lineages (namely BA.2 (Omicron 21L), B.1.160 (Marseille-4), BA.1.1 (Omicron 21K), AY.43 (Delta 21J), and BA.5.1 (Omicron 22B)) harbored >50% of all 220 “hyperfertile” nucleotide mutations. Among the 220 “hyperfertile” mutations, five were mainly found in the B.1.160 (Marseille-4), Delta 21J, and BA.1 (Omicron 21K) or BA.2 (Omicron 21L) variants. One mutation (A18163G / I1566V) was mainly found in the BA.1 (Omicron 21K) and BA.2 (Omicron 21L) variants. One mutation (C19220U / A1918V) was mainly found in the Delta 21J and the BA.2 (Omicron 21L) variants. One mutation (C18744U / Y6160Del) was essentially found in the Delta 21J variant. One mutation (C18877U / Syn.) was mainly found in the B.1.160 (Marseille-4) lineage. Finally, one mutation (C19524U / Syn.) was essentially found in the Delta 21J variant.

### Frequencies of mutations according to SARS-CoV-2 genes and gene categories

The mean (±standard deviation) number of nucleotide mutations per gene was 895±947 (range: 140–4,198; median: 570) (Table 2; Supplementary Figure S5). When taking into account the gene length, the mean number of mutations per 1,000 nucleotides per gene was 825±346 (537–1,852; median: 643) (Figure 3). Also, the mean number of mutations per 1,000 nucleotides per gene was 8±6 (0–22) for “hyperfertile” mutations, 50±27 (20–115) for “fertile” mutations, and 763±316 (497–1,727) for neutral/deleterious mutations, differences between these categories of mutations being statistically significant (p<10^-6^, Welch’s analysis of variance).

**Figure 3.**
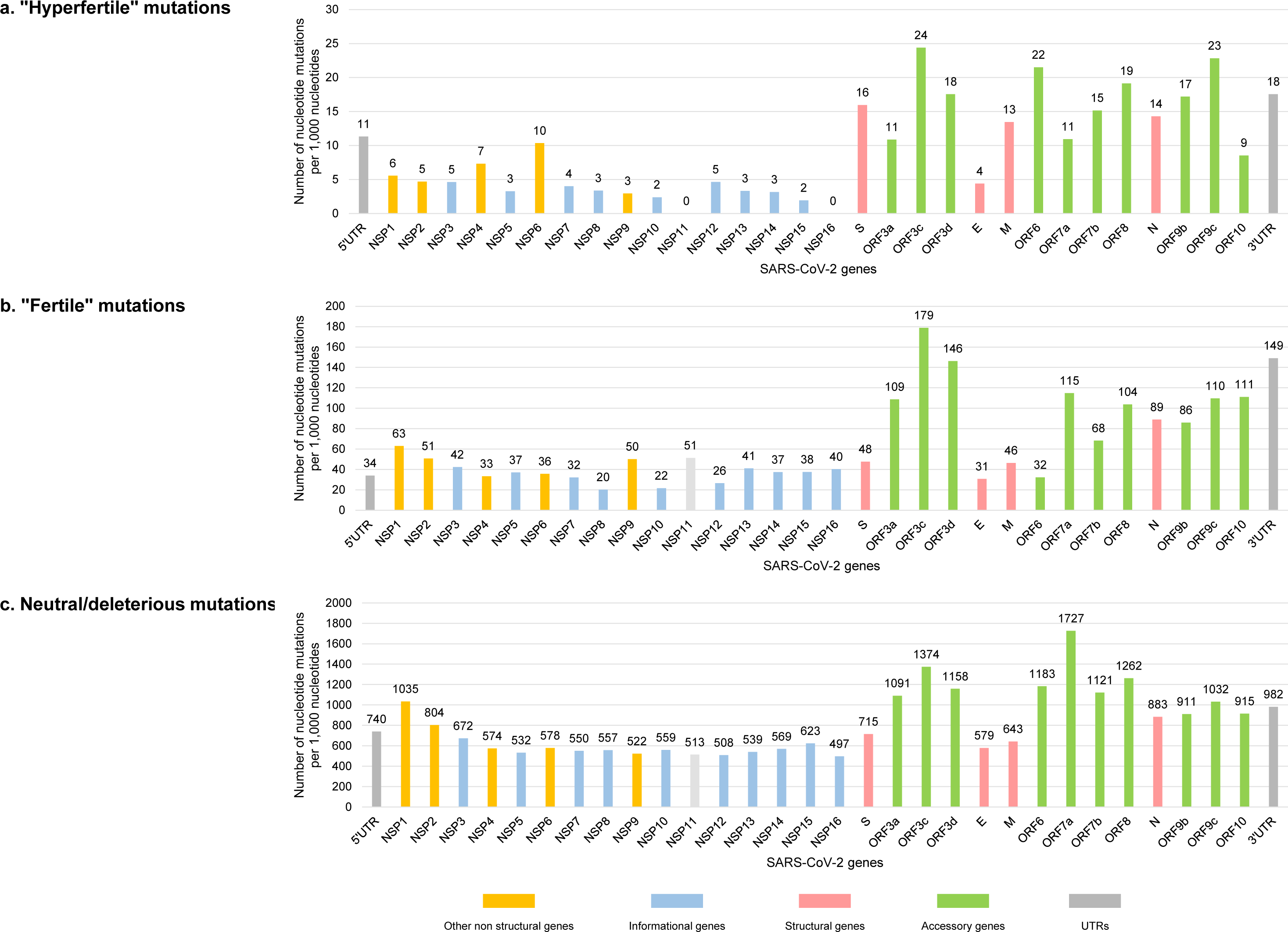
Frequencies of “hyperfertile”, “fertile”, and neutral or deleterious nucleotide mutations according to SARS-CoV-2 genes. Frequencies are mentioned as number of mutations per 1,000 nucleotides per gene. Genes were classified into informational, structural, accessory, and other non-structural genes. E, envelope; M, membrane; N, nucleocapsid; S, spike; UTR, untranslated region.

**Table 2.** Frequencies of nucleotide mutations per SARS-CoV-2 gene or genome region.

| Genes | Nucleotide size | Number of mutations per gene | Number of “hyperfertile” mutations per gene | Number of “fertile” mutations per gene | Number of neutral/deleterious mutations per gene | Number of mutations per 1,000 nucleotides | Number of “hyperfertile” mutations per 1,000 nucleotides | Number of “fertile” mutations per 1,000 nucleotides | Number of neutral/deleterious mutations per 1,000 nucleotides |
| --- | --- | --- | --- | --- | --- | --- | --- | --- | --- |
| 5'UTR | 265 | 1 | 3 | 9 | 196 | 785 | 11 | 34 | 740 |
| NSP1 | 540 | 596 | 3 | 34 | 559 | 1,104 | 6 | 63 | 1,035 |
| NSP2 | 1,914 | 1,645 | 9 | 97 | 1,539 | 859 | 5 | 51 | 804 |
| NSP3 | 5,835 | 4,198 | 27 | 247 | 3,924 | 719 | 5 | 42 | 672 |
| NSP4 | 1,500 | 922 | 11 | 50 | 861 | 615 | 7 | 33 | 574 |
| NSP5 | 918 | 525 | 3 | 34 | 488 | 572 | 3 | 37 | 532 |
| NSP6 | 870 | 543 | 9 | 31 | 503 | 624 | 10 | 36 | 578 |
| NSP7 | 249 | 146 | 1 | 8 | 137 | 586 | 4 | 32 | 550 |
| NSP8 | 594 | 345 | 2 | 12 | 331 | 581 | 3 | 20 | 557 |
| NSP9 | 339 | 195 | 1 | 17 | 177 | 575 | 3 | 50 | 522 |
| NSP10 | 417 | 243 | 1 | 9 | 233 | 583 | 2 | 22 | 559 |
| NSP12 | 39 | 22 | 0 | 2 | 20 | 564 | 0 | 51 | 513 |
| NSP13 | 2,795 | 1,530 | 13 | 74 | 1,421 | 547 | 5 | 26 | 508 |
| NSP14 | 1,803 | 1,052 | 6 | 74 | 972 | 583 | 3 | 41 | 539 |
| NSP15 | 1,581 | 963 | 5 | 59 | 899 | 609 | 3 | 37 | 569 |
| NSP16 | 1,038 | 688 | 2 | 39 | 647 | 663 | 2 | 38 | 623 |
| S | 894 | 480 | 0 | 36 | 444 | 537 | 0 | 40 | 497 |
| ORF3a | 3,822 | 2,974 | 61 | 182 | 2,731 | 778 | 16 | 48 | 715 |
| ORF3c | 828 | 1,002 | 9 | 90 | 903 | 1,210 | 11 | 109 | 1,091 |
| ORF3d | 123 | 594 | 3 | 22 | 169 | 4,829 | 24 | 179 | 1,374 |
| E | 171 | 226 | 3 | 25 | 198 | 1,322 | 18 | 146 | 1,158 |
| M | 228 | 140 | 1 | 7 | 132 | 614 | 4 | 31 | 579 |
| ORF6 | 669 | 470 | 9 | 31 | 430 | 703 | 13 | 46 | 643 |
| ORF7a | 186 | 230 | 4 | 6 | 220 | 1,237 | 22 | 32 | 1,183 |
| ORF7b | 366 | 678 | 4 | 42 | 632 | 1,852 | 11 | 115 | 1,727 |
| ORF8 | 132 | 169 | 2 | 9 | 148 | 1,280 | 15 | 68 | 1,121 |
| N | 366 | 507 | 7 | 38 | 462 | 1,385 | 19 | 104 | 1,262 |
| ORF9b | 1,260 | 1,243 | 18 | 112 | 1,113 | 987 | 14 | 89 | 883 |
| ORF9c | 291 | 295 | 5 | 25 | 265 | 1,014 | 17 | 86 | 911 |
| ORF10 | 219 | 255 | 5 | 24 | 226 | 1,164 | 23 | 110 | 1,032 |
| 3'UTR | 117 | 121 | 1 | 13 | 107 | 1,034 | 9 | 111 | 915 |

E, envelope; M, membrane, N, nucleocapsid; S, spike

Strikingly, the number of “hyperfertile” mutations per 1,000 nucleotides was statistically higher for the accessory genes compared to the informational genes (14 versus 4; 3.7-fold difference; p= 0.0037, Welch Two Sample t-test) (Table 2; Figure 3). Particularly, “hyperfertile” mutations were 4.1-fold, significantly more frequent in ORF8 than in NSP12, which encodes the RNA-dependent RNA polymerase, while they were 6.1-fold more frequent in ORF8 than in NSP14, which encodes the 3’-5’-exonuclease with proofreading activity. The frequency of “hyperfertile” mutations was also significantly higher, 4.0-fold, in structural genes (encoding the spike, membrane, envelope, and nucleocapsid proteins) than in informational genes (15 versus 4 mutations per 1,000 nucleotides). Regarding “fertile” mutations, they were 2.7-fold more frequent in accessory genes that in informational genes (p= 0.028, Welch Two Sample t-test) (Table 2; Figure 3). Also, they were 3.9-fold more frequent in ORF8 than in NSP12, and 2.8-fold more frequent in ORF8 than in NSP14. Finally, neutral or deleterious mutations were 2.1-fold more frequent in accessory genes that in informational genes, 2.5-fold more frequent in ORF8 than in NSP12, and 2.2-fold more frequent in ORF8 than in NSP14 (Table 2).

### Frequencies and localizations in SARS-2 genomes of nucleotide mutations generating stop codons

Stop codons were present at 97 different codons overall. Nine genes were found to harbor at least one mutation generating a stop codon. These genes were by order of stop codon frequency per 100 codons: ORF7a (n= 25 stop codons per 100 codons), ORF8 (n= 21), ORF7b (n= 20), ORF9b (n= 12), ORF6 (n= 10), ORF3a (n= 3), M (n= 1), S (n= 0.2), and ORF1b (n= 0.04) (Figure 4). Therefore, these stop codons were almost only harbored in SARS-CoV-2 genomes by accessory genes, and exceptionally so in structural and other non-structural genes.

**Figure 4.**
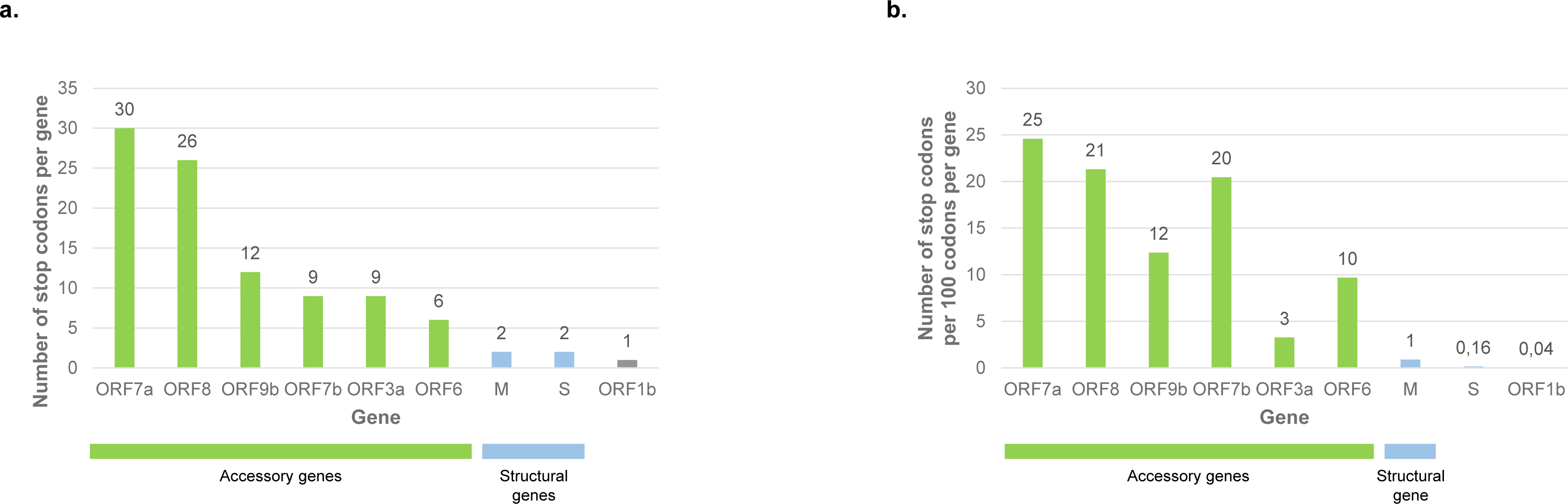
Number of stop codons per gene (a) and per 100 codons per gene (b) Genes were classified into accessory and structural genes. M, membrane; S, spike

## DISCUSSION

SARS-CoV-2 provided an exceptional opportunity to observe the emergence and development of a viral pandemic for which the number of genomic sequences obtained at the different phases of the virus spreading has never reached such a high level, with around 15 million genomes overall in the GISAID sequence bank (https://gisaid.org/; Elbe *et al*., 2017). In our centre alone, we were able to sequence more than 60,000 complete genomes, which we have analyzed here. These genomic data are very important because they demonstrate the extent to which a new infectious disease presents a completely unpredictable evolution, since we cannot predict what changes will appear during the pandemic which may favor the genetic diversification of the virus or not. Moreover, it was not clear at the beginning of the pandemic that there would be animal foci that would possibly also play a role in the appearance of rapidly spreading variants (Qiu *et al*., 2023), while it appears likely currently that mustelids and murids played a role in the pandemic (Reggiani *et al*., 2022). The regular appearance of about one mutation every 15 days in SARS-CoV-2 genomes led after three years of evolution to reach nearly 120 mutations per genomes compared to the first sequenced genome originating from Wuhan (https://nextstrain.org/; Markov *et al*., 2023). Under these conditions, we felt it was important to try to correlate the existence of these mutations with the different epidemic episodes of the virus.

As we have been able to examine it here in Marseille, it seems that the first stage that saw the disease change from being an epidemic in China confined to the province of Hubei to being a pandemic, occurred in Italy with two major phenomena. On the one hand, viruses emerged carrying a mutation (P323L) in the RNA-dependent RNA polymerase (Goldswain *et al*., 2023) which led to a considerable increase in the number of errors during genome replication and therefore in the variability of this virus by increasing the number of mutants in a range that has been speculated to be at least three times higher (Pachetti *et al*., 2020a; Yin *et al*., 2023). This level of random mutation capacity, which fits the Court Jester theory (Benton, 2009), was unanticipated, as other coronaviruses do not show mutation rates of this nature (Worobey *et al*., 2020; Rausch *et al*., 2020). In practice, this rate has been associated with the appearance of one fixed mutation on average every 15 days, and thus the viral genomes currently obtained present more than 100 different mutations relatively to that of the virus that emerged in Wuhan, China, in December 2019. This was also the case for viruses associated with a new epidemic outbreak in China after 33 months of epidemic control (Pan *et al*., 2023). On the other hand, at the same time as this mutation allowed the considerable increase in errors and therefore random mutations, another mutation (D614G) appeared in the spike protein responsible for interaction with the cellular receptor angiotensin-converting enzyme 2 allowing SARS-CoV-2 to enter cells, that promoted infectivity, transmission and probably disease severity (Korber *et al*., 2020). The combination of these two mutations is likely to have been what allowed the virus to trigger a pandemic which has been uncontrolled to date.

In this study, we wanted to quantify the mutations and their effects on the fitness of SARS-CoV-2. Thus, we have defined “hyperfertile” and “fertile” mutations. The most fertile of all these mutations are the two that we have just described in RNA polymerase and spike. A mutation was considered as “hyperfertile” when at least 835 offspring followed its appearance. The selection of mutations during the evolution of the virus appears particularly interesting. First, apart from the key mutation P323L that appeared in RNA polymerase, an informational gene, very few “hyperfertile”/”fertile” mutations appeared in such informational genes that are essential for viral replication. This reasonably reflects that the viral informational machinery needs to maintain its integrity to allow the virus to replicate. Second, many mutations have been observed in the accessory genes. In these genes, we can note three types of phenomena: “hyperfertile” mutations, “fertile” mutations, and neutral/deleterious mutations and, in a way that was extremely surprising, a considerable number of stop codons. For some genes, these stop codons may testify to their limited use in the fitness of the virus, since these mutations have been selected. In contrast, in ORF8, the stop codons, regardless of the site where they appeared, are associated with a resumption of fitness, which is part of the elements that seemed unpredictable (Colson *et al*., 2023). Thus, on at least three occasions, the appearance of a stop codon initiated an epidemic episode, in the cases of the Marseille-4b (Colson *et al*. 2023) and B.1.1.7 (Alpha) (Volz *et al*., 2021) variants and, very recently, of the Omicron XBB.1.5 recombinant (Uriu *et al*., 2023). These are independent phenomena and, therefore, we can consider that ORF8 is not only not essential, but is in fact harmful to the virus at this stage of the epidemic, perhaps due to cross-reactivities with circulating viruses that would have led to a negative selection pressure on its epitopes (Hachim *et al*., 2020; Colson *et al*., 2023). This could be an explanation for the apparent immunity of the youngest individuals at the start of the pandemic, which could be the trace of cross-infections with other viruses (Colson *et al*., 2020; Jungreis *et al*., 2021). It should be noted that among the SARS-CoV-2 genes, ORF8 is the one that very significantly accumulates the largest number of mutations, even outside of the mutations generating stop codons. This shows that it is, most of the time, useless and often deleterious for the development of epidemics. It is possible that this phenomenon of knocking out ORF8 makes it possible to understand that the specialization of a virus for a new host is associated with a loss of genes that are not only useless, as is the case of other accessory genes, since their disappearance does not prevent the continuation of epidemics, but that could hinder the viral replication due to immunological cross-reactivities. Thus, the general mechanism of pathogenicity for hyper-specialized infectious agents for a given animal group is, probably in most cases, in the same pathway, i.e., through genomic reduction with conservation only of what is used and particularly what is essential, corresponding to the paradigm of “use it or lose it” (Moran *et al*., 2002; Colson *et al*., 2023).

## Author Contributions

D.R. and P.C. designed the study. P.C., H.C., J.D., AL., J.F., C.D., and D.R. provided materials, data or analysis tools. All authors analyzed the data. D.R. and P.C. wrote the first draft of the manuscript. All authors reviewed and approved the final manuscript.

## Funding

This study was supported by the French Government under the “Investments for the Future” program managed by the National Agency for Research (ANR) (Méditerranée-Infection 10-IAHU-03); by the Région Provence Alpes Côte d’Azur and European funding FEDER PRIMMI (Fonds Européen de Développement Régional-Plateformes de Recherche et d’Innovation Mutualisées Méditerranée Infection) (FEDER PA 0000320 PRIMMI); and by the French Ministry of Higher Education, Research and Innovation (Ministère de l’Enseignement supérieur, de la Recherche et de l’Innovation) and the French Ministry of Solidarity and Health (Ministère des Solidarités et de la Santé) in the framework of the Consortium for Surveillance and Research of EMERgent Pathogen Infections via Microbial GENomics (EMERGEN) (https://www.santepubliquefrance.fr/dossiers/coronavirus-covid-19/consortium-emergen).

**Table 3.** Location and frequency of stop codons.

| Gene : Codon | Frequency |
| --- | --- |
| ORF8: Q27* | 3,930 |
| ORF8: K68* | 1,304 |
| ORF8: E64* | 998 |
| ORF7a: Q62* | 172 |
| ORF7a: G38* | 109 |
| ORF8: Q18* | 86 |
| ORF7a: E121* | 82 |
| ORF8: E106* | 73 |
| ORF7b: E3* | 55 |
| ORF7b: E39* | 50 |
| ORF8: G8* | 47 |
| ORF8: E19* | 43 |
| ORF6: E54* | 36 |
| ORF7a: E95* | 30 |
| ORF7a: *122R | 2 |
| ORF3a: G254* | 27 |
| ORF7a: *122L | 8 |
| ORF8: E92* | 21 |
| ORF8: E110* | 21 |
| ORF7a: Q94* | 20 |
| ORF7b: *44Y | 2 |
| ORF8: L60* | 20 |
| ORF7a: Q90* | 19 |
| ORF7a: E91* | 18 |
| ORF7a: *122G | 6 |
| ORF7b: *44- | 18 |
| ORF8: G50* | 16 |
| ORF7a: L77* | 15 |
| ORF9b: K67* | 15 |
| ORF7a: E92* | 14 |
| ORF3a: E261* | 12 |
| ORF7a: *122- | 12 |
| ORF8: W45* | 12 |
| ORF8: Q72* | 12 |
| ORF8: E59* | 11 |
| ORF3a: Q245* | 10 |
| ORF6: *62- | 8 |
| ORF7a: Q76* | 8 |
| ORF8: Q23* | 8 |
| ORF8: Q29* | 8 |
| ORF9b: *98S | 1 |
| ORF6: Q8* | 7 |
| ORF7a: E41* | 7 |
| ORF7a: K117* | 7 |
| ORF9b: Q77* | 7 |
| ORF9b: *98R | 3 |
| ORF6: E55* | 6 |
| ORF8: L95* | 6 |

| Gene: Codon | Frequency |
| --- | --- |
| ORF7b: E33* | 5 |
| ORF9b: Q34* | 5 |
| ORF8: Y31* | 4 |
| ORF9b: E65* | 4 |
| ORF9b: E86* | 4 |
| ORF9b: *98L | 4 |
| ORF7a: C35* | 3 |
| ORF7a: K85* | 3 |
| ORF7b: W29* | 3 |
| ORF9b: Q18* | 3 |
| ORF9b: Q20* | 3 |
| ORF9b: E27* | 3 |
| ORF3a: G44* | 2 |
| ORF3a: E241* | 2 |
| ORF6: W27* | 2 |
| ORF7a: Q21* | 2 |
| ORF7a: R25* | 2 |
| ORF7a: E33* | 2 |
| ORF7a: Y40* | 2 |
| ORF7a: Y97* | 2 |
| ORF7a: K119* | 2 |
| ORF8: S69* | 2 |
| ORF8: Q91* | 2 |
| ORF9b: K4* | 2 |
| M: Y204* | 1 |
| M: *223- | 1 |
| ORF1b:<br>Y1944* | 1 |
| ORF3a: Q38* | 1 |
| ORF3a: E181* | 1 |
| ORF3a: E194* | 1 |
| ORF3a: E242* | 1 |
| ORF6: Q56* | 1 |
| ORF7a: L7* | 1 |
| ORF7a: L31* | 1 |
| ORF7a: C58* | 1 |
| ORF7a: C67* | 1 |
| ORF7a: Y75* | 1 |
| ORF7a: R89* | 1 |
| ORF7b: L11* | 1 |
| ORF7b: L14* | 1 |
| ORF7b: F19* | 1 |
| ORF8: H17* | 1 |
| ORF8: Y46* | 1 |
| ORF8: K53* | 1 |
| ORF8: S54* | 1 |
| ORF8: Y79* | 1 |
| ORF8: *122L | 1 |
| S: K202* | 1 |
| S: E1262* | 1 |

## Acknowledgments

Part of the SARS-CoV-2 genomes obtained since 2021 have been sequenced in the framework of the EMERGEN program (https://www.santepubliquefrance.fr/dossiers/coronavirus-covid-19/consortium-emergen) from respiratory samples originating from other laboratories than ours, which we thank.

## Ethical statement

SARS-CoV-2 genomes were obtained through genomic surveillance, as recommended by the French government. SARS-CoV-2 genome sequencing had been approved by the Ethical Committee of the Méditerranée Infection institute under reference No. 2020-016-3. Access to the patients’ biological and registry data issued from the hospital information system was approved by the data protection committee of the Assistance Publique-Hopitaux de Marseille (APHM) and was recorded in the European General Data Protection Regulation registry under number RGPD/APHM 2019-73; the latter authorization allowed us to access the patients’ data retrospectively, according to the French law on “Règlement Général sur la Protection des Données (RGPD)”.

## Data Availability Statement

SARS-CoV-2 genomes are available in the GenBank sequence database (https://www.ncbi.nlm.nih.gov/genbank/, Sayers *et al*., 2023), the GISAID sequence database (https://gisaid.org/, Elbe *et al*., 2017), or on the website of the Méditerranée Infection university hospital institute (https://www.mediterranee-infection.com/acces-ressources/donnees-pour-articles/).

