## Supplementary material for "Role of SARS-CoV-2 mutations in the evolution of the COVID-19 pandemic"

### FOR ARTICLE:

**Full-length title: Role of SARS-CoV-2 mutations in the evolution of the COVID-19 pandemic**

**Affiliations:** [1]IHU Méditerranée Infection, 19–21 Boulevard Jean Moulin, 13005 Marseille, France; [2]Aix-Marseille Université, Institut de Recherche pour le Développement (IRD), Microbes Evolution Phylogeny and Infections (MEPHI), 27 Boulevard Jean Moulin, 13005 Marseille, France; [3]Assistance Publique-Hôpitaux de Marseille (AP-HM), 264 Rue Saint-Pierre, 13005 Marseille, France; [4]Aix-Marseille Université, Institut de Recherche pour le Développement (IRD), Vecteurs - Infections Tropicales et Méditerranéennes (VITROME), 27 Boulevard Jean Moulin, 13005 Marseille, France; [5] French Armed Forces Center for Epidemiology and Public Health (CESPA), Camp de Sainte Marthe, Marseille, France; [6]Department of Biological Sciences, Centre National de la Recherche Scientifique, Centre National de la Recherche Scientifique (CNRS)-SNC5039, Marseille, France; [7] INSERM UMR_S 1072, Aix-Marseille Université, Marseille, France

#### SUPPLEMENTARY FIGURES

**Supplementary Figure S1. Daily numbers of new SARS-CoV-2 diagnoses (a) and distribution of the most predominant SARS-CoV-2 variants over the whole pandemic period based on genomes described in the present study (b)**

**Suppl. Fig. S1**

**a.**

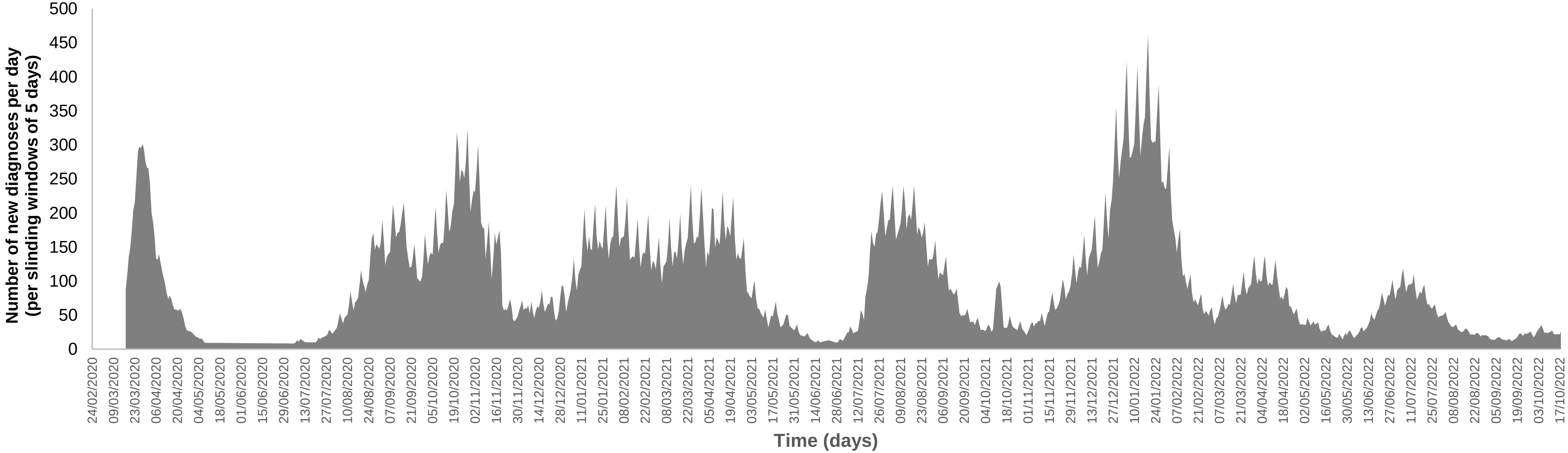


**b.**

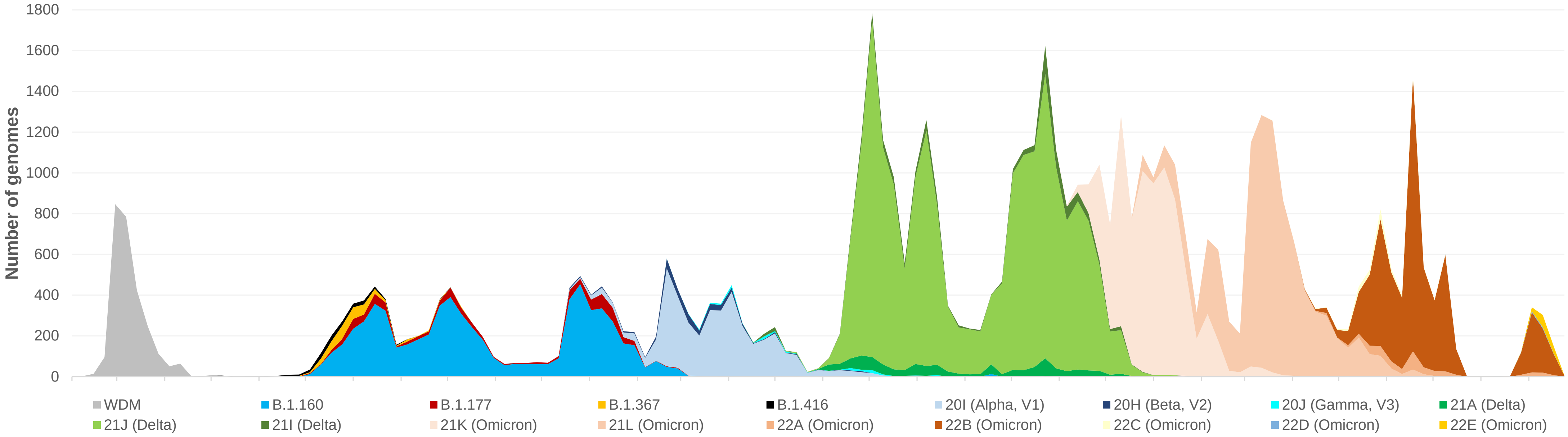

**Supplementary Figure S2. Evolution over time of the number of any nucleotide mutations (a) and of "hyperfertile" nucleotide mutations (b) per genome**

**Supplementary Fig. S2**

**a.**

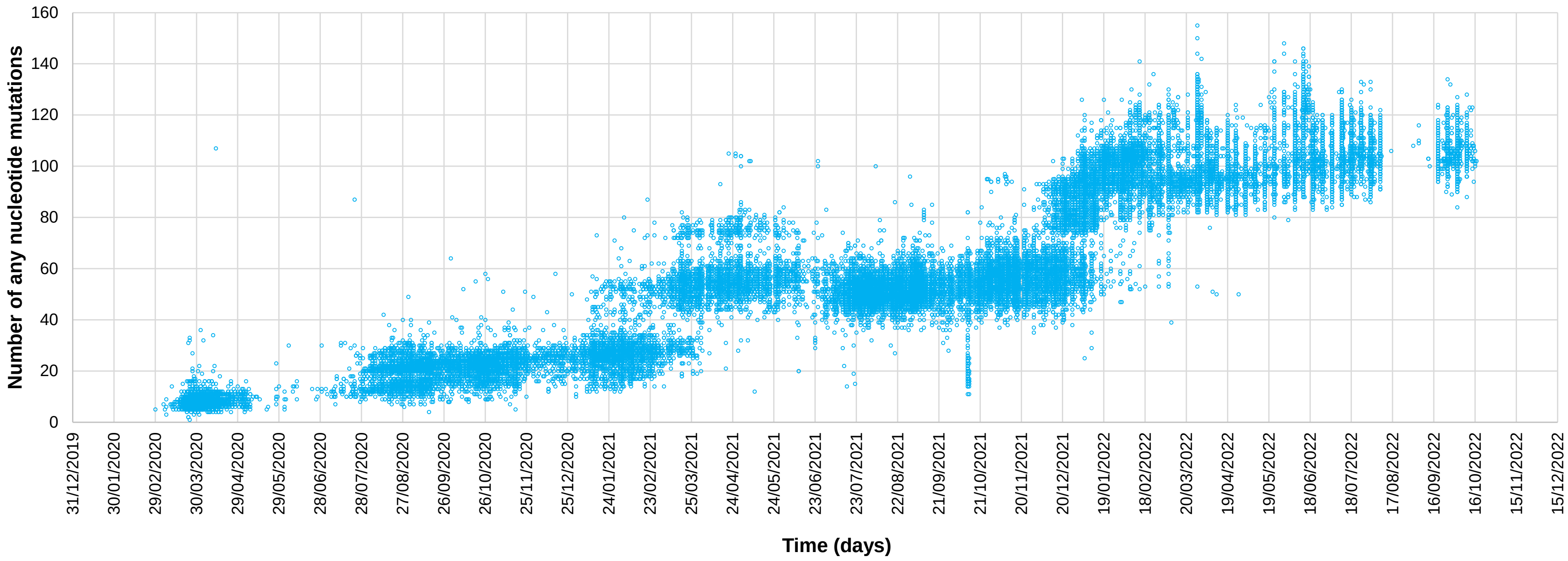


**b.**

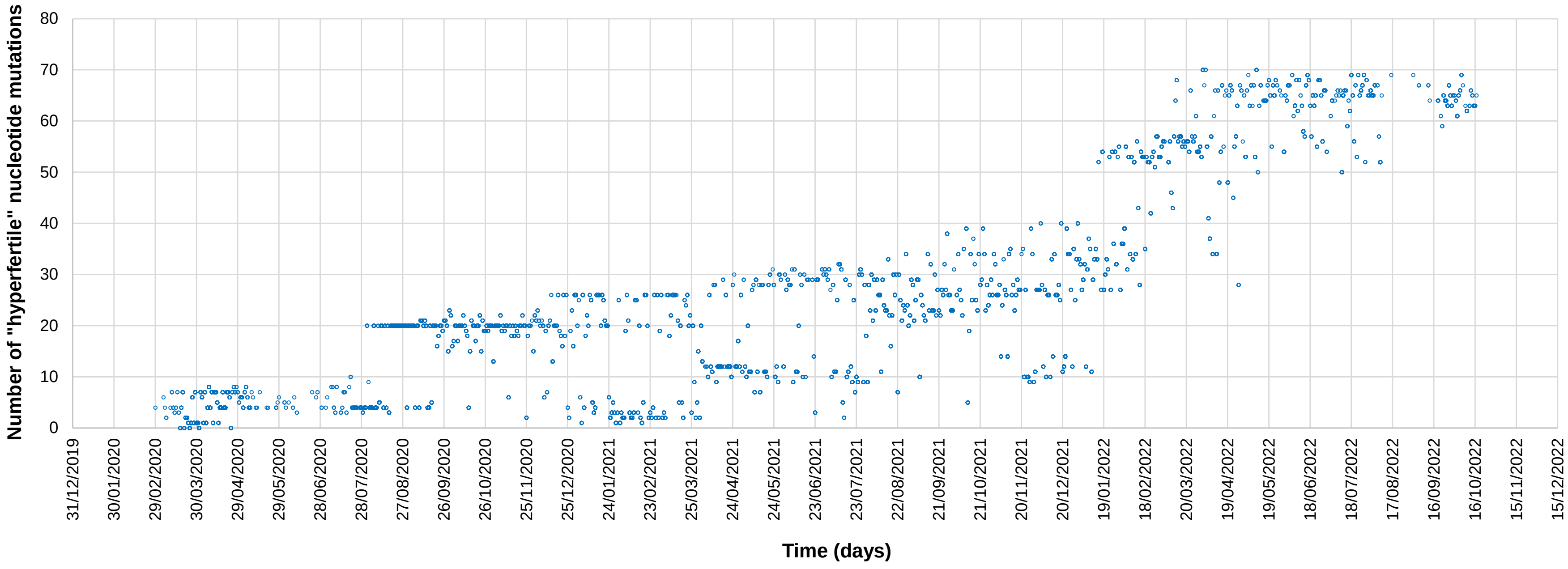

**Supplementary Figure S3. Distribution of the frequencies of nucleotide (a, c) and amino acid (b, d) mutations, and of mutations generating stop codons among the 61,397 SARS-CoV-2 genomes obtained in our institute**

**Suppl. Fig. S3**

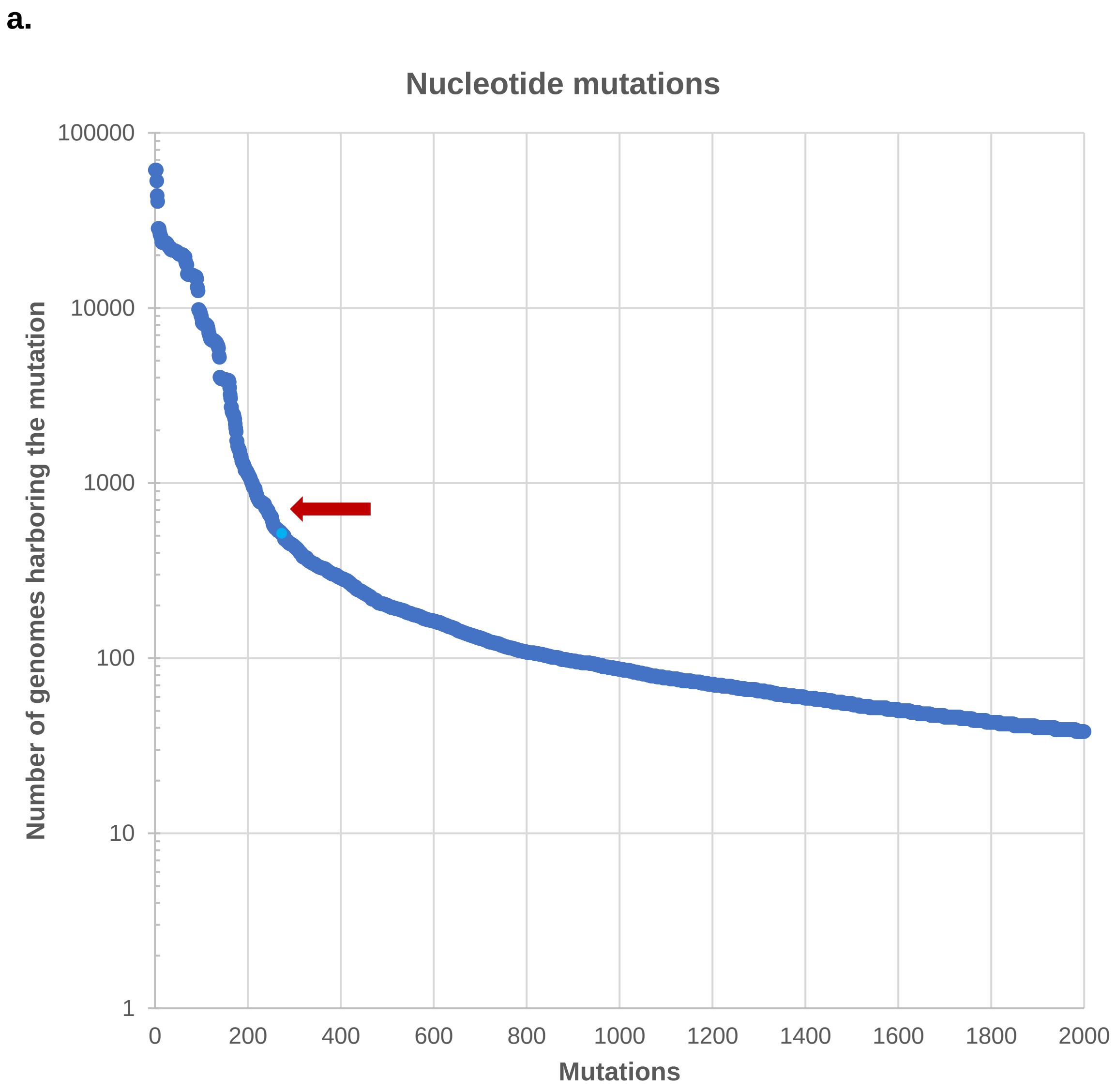
a.
Nucleotide mutations
Number of genomes harboring the mutation
100000
10000
1000
100
10
1
0
200
400
600
800
1000
1200
1400
1600
1800
2000
Mutations

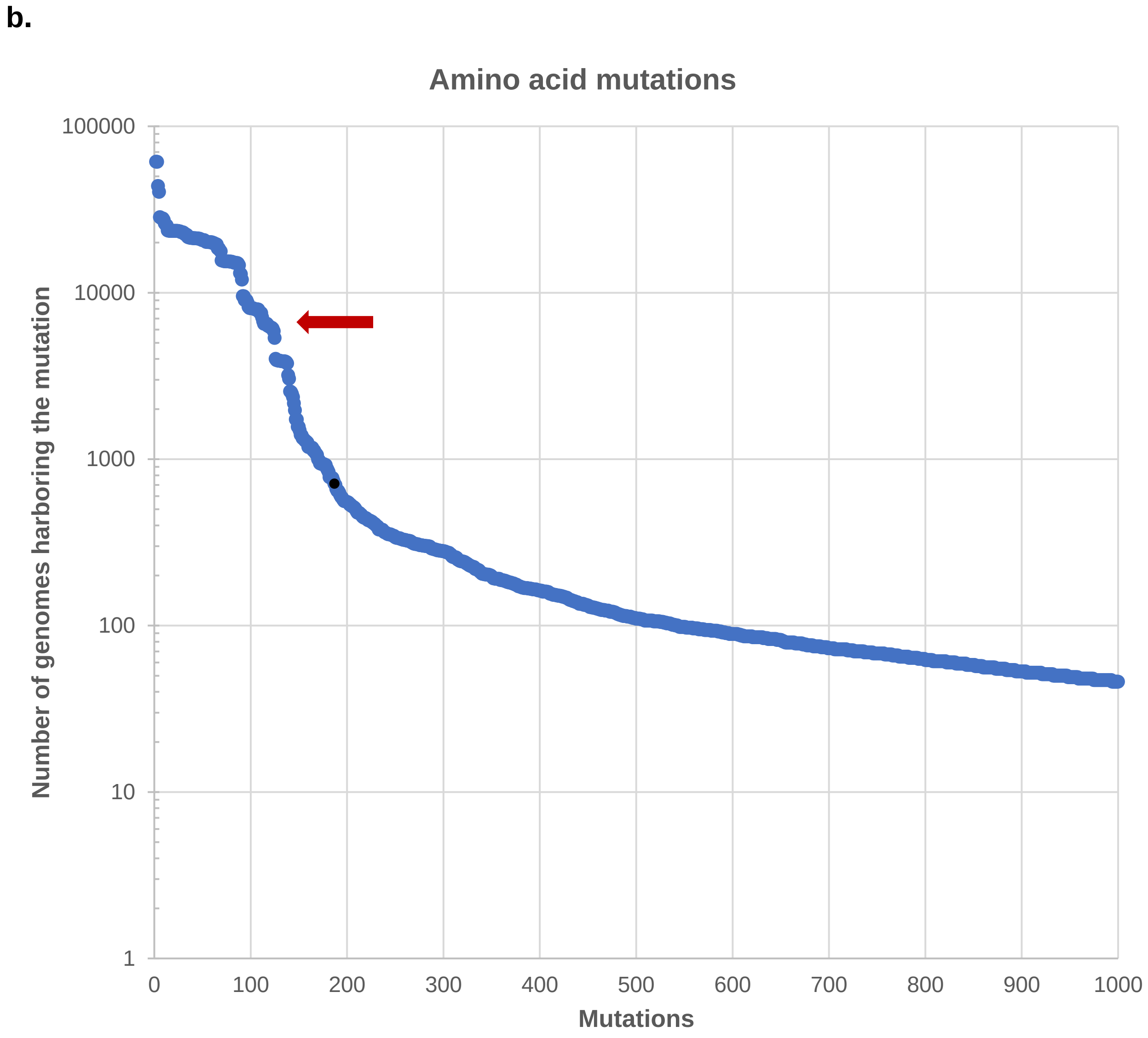
b.
Amino acid mutations
Number of genomes harboring the mutation
100000
10000
1000
100
10
1
0
100
200
300
400
500
600
700
800
900
1000
Mutations

Suppl. Fig. S3

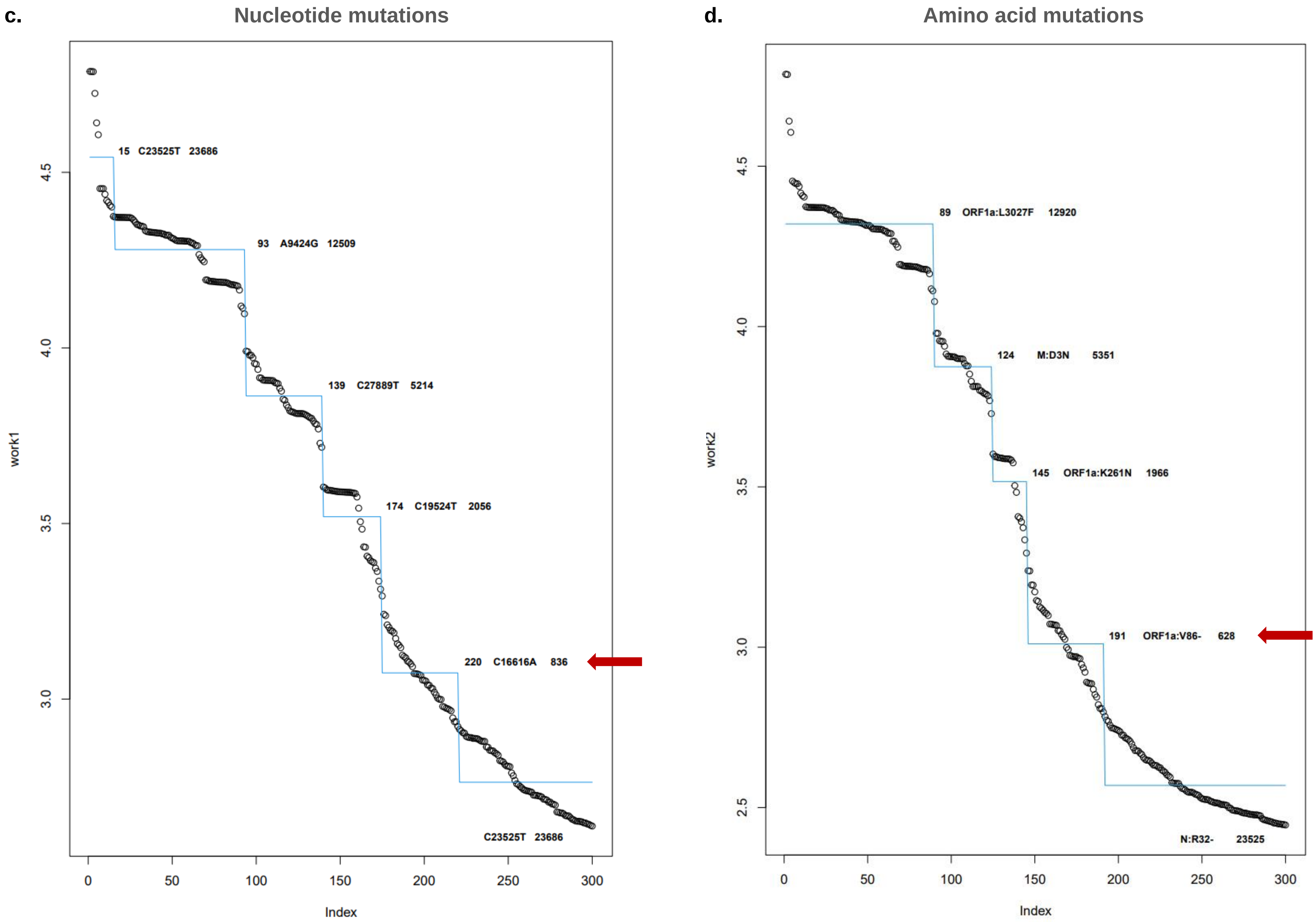
c.
Nucleotide mutations
work1
Index
15 C23525T 23686
93 A9424G 12509
139 C27889T 5214
174 C19524T 2056
220 C16616A 836
C23525T 23686
d.
Amino acid mutations
work2
Index
89 ORF1a:L3027F 12920
124 M:D3N 5351
145 ORF1a:K261N 1966
191 ORF1a:V86- 628
N:R32- 23525

**Suppl. Fig. S3**

**e.**

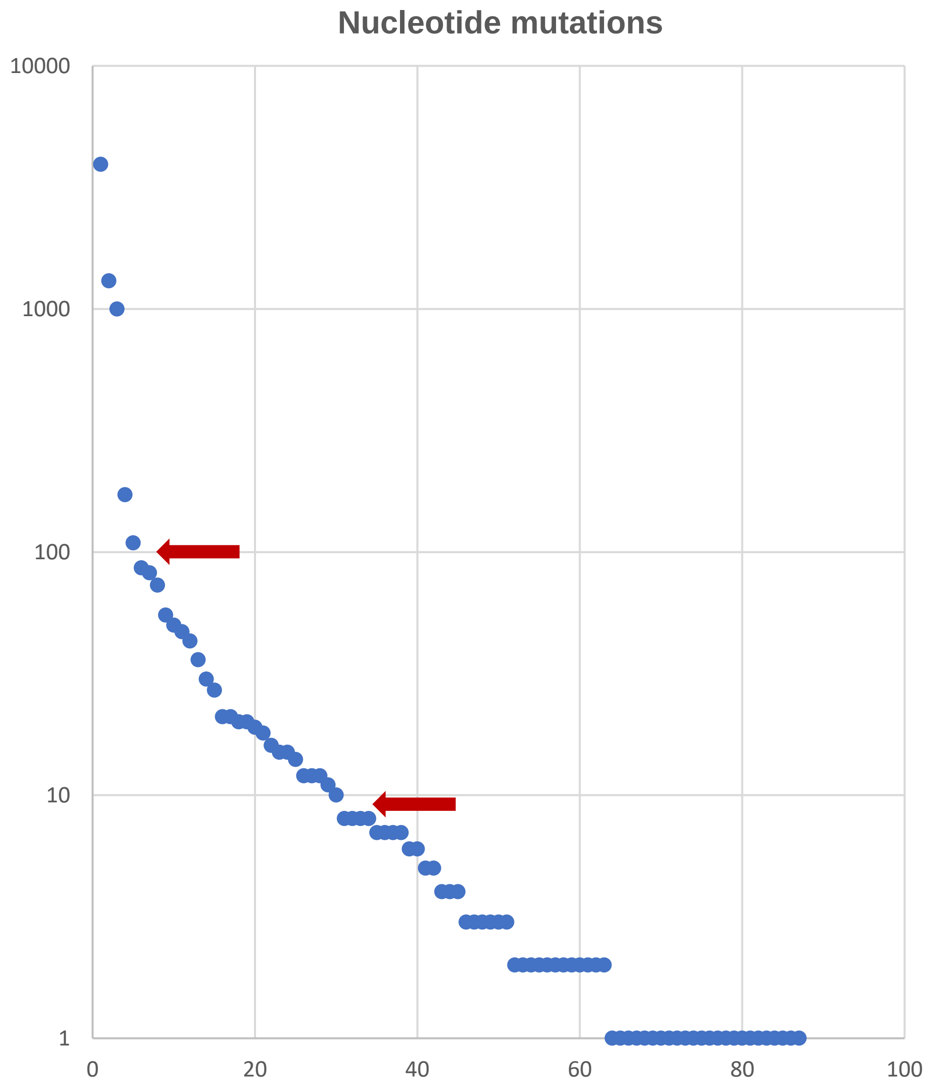

**Supplementary Figure S4. Structures of the Wuhan-Hu-1 SARS-CoV-2 RNA-dependent RNA polymerase (a) and spike (c) proteins and of their mutant forms incorporating the P323L (b) and D614G (d) amino acid substitutions, respectively**

a, b: P323L induces "long range" conformational effect that modifies the interaction of the RNA polymerase with the primer and the RNA template (Pachetti *et al.*, 2020a). C, d: D614G enables facilitation of the conformational change that unmasks the cellular receptor (ACE-2) in the spike trimer (Korber *et al.*, 2020).

RdRp, RNA-dependent RNA polymerase.

**Supplementary Fig. S4**

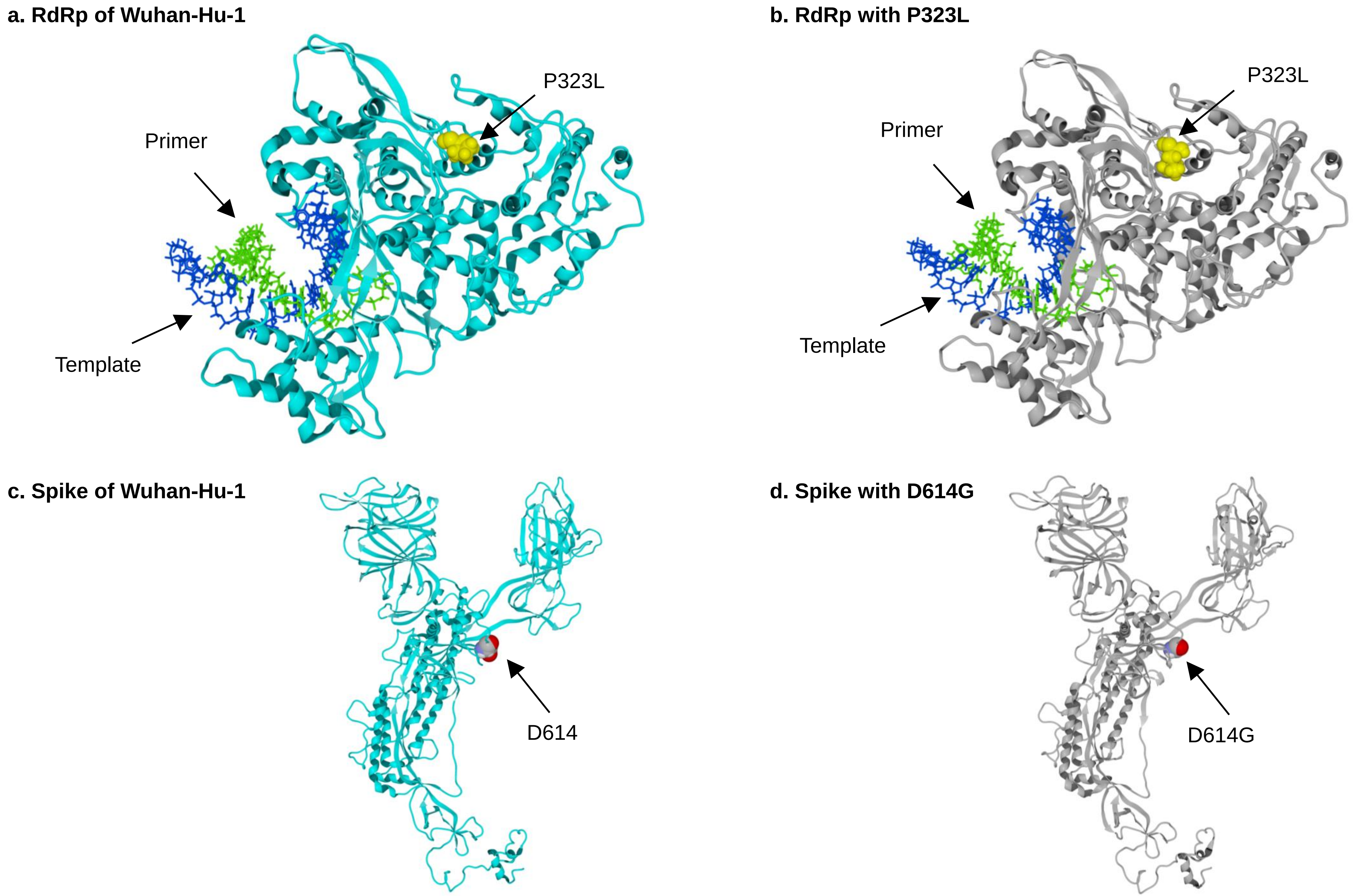

**Supplementary Figure S5. Frequencies of "hyperfertile", "fertile", and neutral or deleterious nucleotide mutations according to SARS-CoV-2 genes**

Genes were classified into informational, structural, accessory, and other non-structural genes. UTR, untranslated region.

#### Supplementary Fig. S5

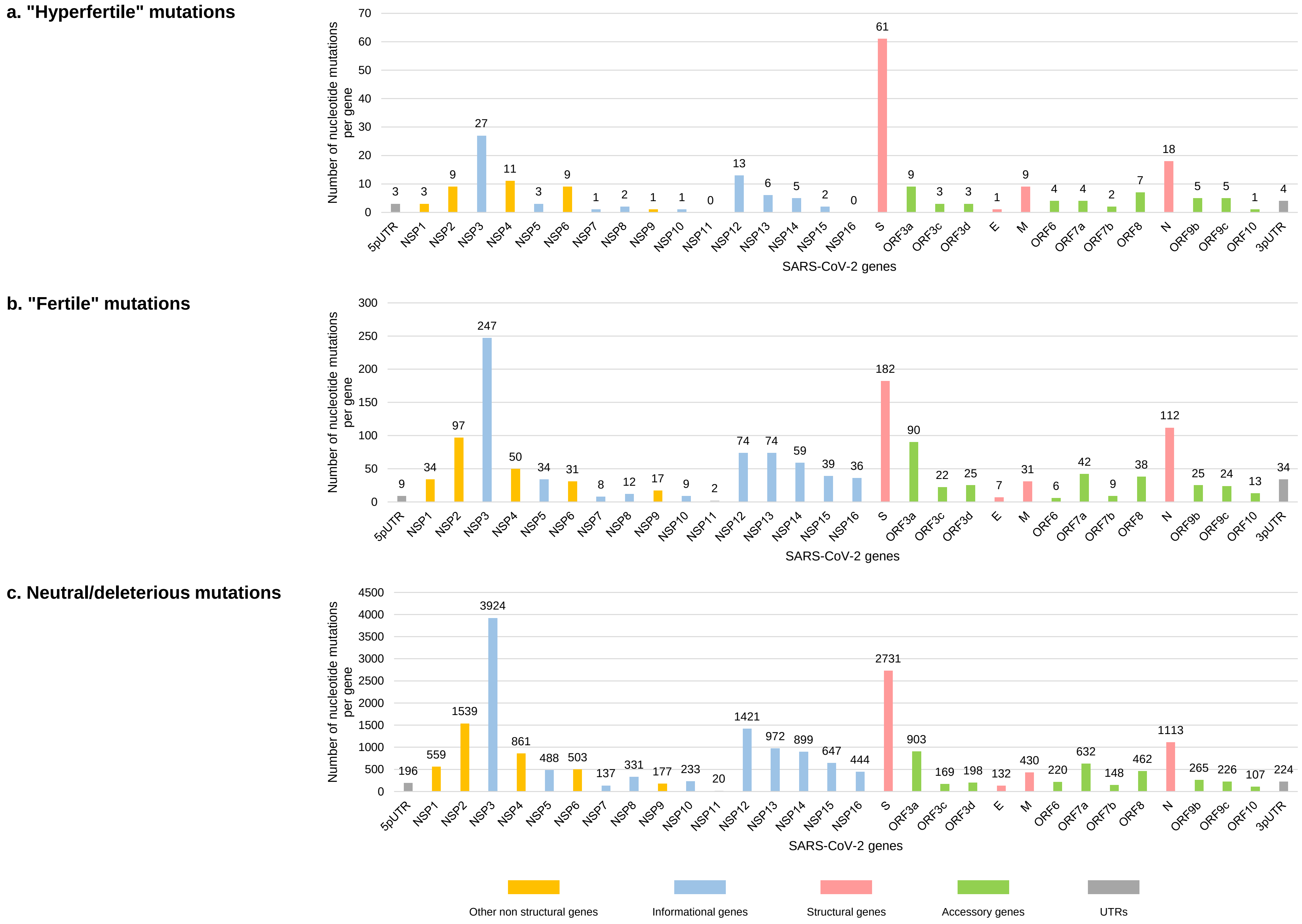

#### SUPPLEMENTARY TABLES

**Supplementary Table S1. List and frequency of the Pangolin lineages and Nextstrain clades into which were classified the 61,397 genomes**

Nextstrain (and WHO) clades (https://nextstrain.org/; Hadfield *et al.*, 2018; https://www.who.int/activities/tracking-SARS-CoV-2-variants) and Pangolin lineages (https://cov-lineages.org/resources/pangolin.html; Rambaut *et al.*, 2020) were determined using the Nextclade tool (https://clades.nextstrain.org/; Aksamentov *et al.*, 2021).

| Pangolin lineage | Nextstrain (WHO) clades | Frequency |
|---|---|---|
| AY.43 | 21J (Delta) | 8992 |
| BA.2 | 21L (Omicron) | 6564 |
| B.1.160 | 20A | 6505 |
| B.1.1.7 | 20I (Alpha, V1) | 3585 |
| B.1 | 20A | 2728 |
| AY.98.1 | 21J (Delta) | 2452 |
| BA.1.1 | 21K (Omicron) | 2298 |
| BA.5.1 | 22B (Omicron) | 1765 |
| BA.1.1.1 | 21K (Omicron) | 1555 |
| AY.42 | 21J (Delta) | 1387 |
| BA.1.18 | 21K (Omicron) | 1211 |
| AY.4 | 21J (Delta) | 1210 |
| BA.1.17 | 21K (Omicron) | 1147 |
| AY.122 | 21J (Delta) | 1007 |
| BA.1 | 21K (Omicron) | 952 |
| AY.122.6 | 21J (Delta) | 948 |
| B.1.617.2 | 21A (Delta) | 883 |
| BA.5.2.1 | 22B (Omicron) | 775 |
| BA.5.2 | 22B (Omicron) | 717 |
| BA.2.9 | 21L (Omicron) | 700 |
| AY.9.2 | 21I (Delta) | 649 |
| AY.125 | 21J (Delta) | 646 |
| B.1.177 | 20E (EU1) | 611 |
| BA.2.9.3 | 21L (Omicron) | 499 |
| BA.1.17.2 | 21K (Omicron) | 411 |
| B.1.1 | 20B | 363 |
| B.1.367 | 20C | 317 |
| B.1.1.269 | 20B | 301 |
| AY.5 | 21J (Delta) | 295 |
| BA.5.1.17 | 22B (Omicron) | 280 |
| BA.2.49 | 21L (Omicron) | 256 |
| AY.109 | 21J (Delta) | 255 |
| AY.34 | 21J (Delta) | 247 |
| BA.5 | 22B (Omicron) | 236 |
| AY.33 | 21J (Delta) | 204 |
| AY.4.2.3 | 21J (Delta) | 203 |
| BA.5.1.22 | 22B (Omicron) | 201 |
| BA.2.3 | 21L (Omicron) | 194 |
| Q.7 | 20I (Alpha, V1) | 175 |
| B.1.1.241 | 20B | 174 |
| BA.5.2.2 | 22B (Omicron) | 165 |
| B.1.351.2 | 20H (Beta, V2) | 164 |
| B.1.221 | 20A | 162 |
| BA.4 | 22A (Omicron) | 162 |
| BA.2.12.1 | 22C (Omicron) | 153 |
| B.1.351 | 20H (Beta, V2) | 152 |
| BA.4.1 | 22A (Omicron) | 152 |

| Pangolin lineage | Nextstrain (WHO) clades | Frequency |
|---|---|---|
| AY.34.1 | 21J (Delta) | 148 |
| B.1.416 | 20A | 136 |
| BE.1.1 | 22B (Omicron) | 133 |
| BA.5.2.20 | 22B (Omicron) | 131 |
| BF.7 | 22B (Omicron) | 128 |
| BE.1 | 22B (Omicron) | 125 |
| AY.5.4 | 21J (Delta) | 124 |
| BA.2.36 | 21L (Omicron) | 121 |
| Q.4 | 20I (Alpha, V1) | 119 |
| BA.5.1.23 | 22B (Omicron) | 112 |
| AY.4.2 | 21J (Delta) | 108 |
| B.1.525 | 21D (Eta) | 106 |
| BA.2.6 | 21L (Omicron) | 104 |
| AY.126 | 21J (Delta) | 103 |
| BA.5.1.10 | 22B (Omicron) | 103 |
| BQ.1.1 | 22E (Omicron) | 99 |
| BA.1.1.14 | 21K (Omicron) | 96 |
| BA.1.15 | 21K (Omicron) | 93 |
| B.1.177.81 | 20E (EU1) | 91 |
| BF.5 | 22B (Omicron) | 90 |
| AY.16.1 | 21A (Delta) | 89 |
| BA.1.13 | 21K (Omicron) | 79 |
| BF.1 | 22B (Omicron) | 77 |
| P.1 | 20J (Gamma, V3) | 73 |
| B.1.258 | 20A | 69 |
| AY.98 | 21J (Delta) | 66 |
| A.21 | 19B | 65 |
| AY.34.2 | 21J (Delta) | 63 |
| BA.4.6 | 22A (Omicron) | 63 |
| AY.46.6 | 21J (Delta) | 61 |
| AY.127 | 21J (Delta) | 59 |
| AY.121 | 21J (Delta) | 58 |
| AY.37 | 21I (Delta) | 56 |
| AY.23 | 21J (Delta) | 48 |
| AY.106 | 21J (Delta) | 47 |
| BA.2.56 | 21L (Omicron) | 45 |
| BF.2 | 22B (Omicron) | 45 |
| BA.5.1.2 | 22B (Omicron) | 45 |
| BA.2.22 | 21L (Omicron) | 43 |
| BA.5.2.3 | 22B (Omicron) | 42 |
| B.1.416.1 | 20A | 41 |
| B.1.356 | 20C | 40 |
| BA.2.18 | 21L (Omicron) | 40 |
| B.1.1.1 | 20D | 39 |
| AY.71 | 21I (Delta) | 39 |
| AY.6 | 21J (Delta) | 39 |
| BA.2.23 | 21L (Omicron) | 38 |

| Pangolin lineage | Nextstrain (WHO) clades | Frequency |
|---|---|---|
| BA.5.5 | 22B (Omicron) | 38 |
| B.1.177.32 | 20E (EU1) | 36 |
| AY.124 | 21J (Delta) | 36 |
| BA.1.15.1 | 21K (Omicron) | 35 |
| AY.120 | 21J (Delta) | 34 |
| P.1.16 | 20J (Gamma, V3) | 33 |
| AY.53 | 21A (Delta) | 32 |
| AY.33.1 | 21J (Delta) | 31 |
| BA.1.21 | 21K (Omicron) | 31 |
| XAZ | recombinant | 31 |
| AY.46.2 | 21J (Delta) | 30 |
| BA.2.65 | 21L (Omicron) | 30 |
| BA.5.1.8 | 22B (Omicron) | 30 |
| BA.1.1.15 | 21K (Omicron) | 29 |
| AY.43.7 | 21J (Delta) | 28 |
| BA.1.14 | 21K (Omicron) | 28 |
| BA.2.1 | 21L (Omicron) | 28 |
| AY.20 | 21J (Delta) | 27 |
| BA.5.1.3 | 22B (Omicron) | 27 |
| AY.92 | 21J (Delta) | 26 |
| BA.5.8 | 22B (Omicron) | 26 |
| AY.112 | 21J (Delta) | 25 |
| AY.36 | 21J (Delta) | 25 |
| AY.73 | 21I (Delta) | 24 |
| AY.72 | 21I (Delta) | 24 |
| AY.118 | 21J (Delta) | 24 |
| BA.5.3 | 22B (Omicron) | 24 |
| B.1.1.297 | 20B | 23 |
| AY.100 | 21J (Delta) | 23 |
| BA.2.9.4 | 21L (Omicron) | 23 |
| BA.5.6 | 22B (Omicron) | 22 |
| BF.10 | 22B (Omicron) | 22 |
| BA.5.9 | 22B (Omicron) | 22 |
| AY.46 | 21J (Delta) | 21 |
| AY.4.2.2 | 21J (Delta) | 21 |
| BA.5.1.24 | 22B (Omicron) | 21 |
| B.1.214.2 | 20A | 19 |
| B.1.1.33 | 20B | 19 |
| B.1.1.318 | 20B | 19 |
| AY.4.4 | 21J (Delta) | 19 |
| BA.1.19 | 21K (Omicron) | 19 |
| BA.5.1.30 | 22B (Omicron) | 19 |
| B.1.147 | 20A | 18 |
| AY.51 | 21A (Delta) | 18 |
| AY.133 | 21I (Delta) | 18 |
| BA.1.16 | 21K (Omicron) | 18 |
| BA.2.10 | 21L (Omicron) | 18 |

| Pangolin lineage | Nextstrain (WHO) clades | Frequency |
|---|---|---|
| BA.4.4 | 22A (Omicron) | 18 |
| BA.5.3.2 | 22B (Omicron) | 18 |
| B.1.640.2 | 20A | 17 |
| B.1.177.73 | 20E (EU1) | 17 |
| AY.7.2 | 21J (Delta) | 17 |
| A.27 | 19B | 16 |
| B.1.389 | 20A | 16 |
| B.1.1.10 | 20B | 16 |
| AY.42.1 | 21J (Delta) | 16 |
| BA.2.3.15 | 21L (Omicron) | 16 |
| BA.2.38 | 21L (Omicron) | 16 |
| BF.28 | 22B (Omicron) | 16 |
| BA.5.3.1 | 22B (Omicron) | 15 |
| BA.5.2.9 | 22B (Omicron) | 15 |
| BG.2 | 22C (Omicron) | 15 |
| BQ.1 | 22E (Omicron) | 15 |
| BA.1.20 | 21K (Omicron) | 14 |
| BA.1.1.11 | 21K (Omicron) | 14 |
| B.1.36 | 20A | 13 |
| B.1.474 | 20A | 13 |
| C.16 | 20D | 13 |
| BA.2.58 | 21L (Omicron) | 13 |
| BA.5.1.5 | 22B (Omicron) | 13 |
| AY.75 | 21I (Delta) | 12 |
| AY.4.7 | 21J (Delta) | 12 |
| AY.119 | 21J (Delta) | 12 |
| AY.123 | 21J (Delta) | 12 |
| B.1.219 | 20A | 11 |
| AY.66 | 21I (Delta) | 11 |
| AY.22 | 21J (Delta) | 11 |
| AY.129 | 21J (Delta) | 11 |
| AY.4.5 | 21J (Delta) | 11 |
| BA.1.4 | 21K (Omicron) | 11 |
| BA.2.52 | 21L (Omicron) | 11 |
| BA.5.3.3 | 22B (Omicron) | 11 |
| A | 19B | 10 |
| B.1.1.189 | 20B | 10 |
| AY.70 | 21I (Delta) | 10 |
| AY.116 | 21J (Delta) | 10 |
| BA.1.8 | 21K (Omicron) | 10 |
| BA.1.1.10 | 21K (Omicron) | 10 |
| BA.2.10.1 | 21L (Omicron) | 10 |
| BA.2.13 | 21L (Omicron) | 10 |
| BA.2.37 | 21L (Omicron) | 10 |
| BA.4.1.1 | 22A (Omicron) | 10 |
| BA.5.2.25 | 22B (Omicron) | 10 |
| BF.7.5 | 22B (Omicron) | 10 |

| Pangolin lineage | Nextstrain (WHO) clades | Frequency |
|---|---|---|
| B.1.620 | 20A | 9 |
| B.1.1.25 | 20B | 9 |
| B.1.621 | 21H (Mu) | 9 |
| AY.46.5 | 21J (Delta) | 9 |
| BA.2.71 | 21L (Omicron) | 9 |
| BE.3 | 22B (Omicron) | 9 |
| BA.5.2.21 | 22B (Omicron) | 9 |
| XZ | recombinant | 9 |
| B | 19A | 8 |
| B.1.640.1 | 20A | 8 |
| B.1.1.371 | 20B | 8 |
| W.1 | 20E (EU1) | 8 |
| AY.9 | 21I (Delta) | 8 |
| AY.9.2.2 | 21I (Delta) | 8 |
| AY.103 | 21J (Delta) | 8 |
| AY.4.2.4 | 21J (Delta) | 8 |
| BA.4.1.10 | 22A (Omicron) | 8 |
| BF.4 | 22B (Omicron) | 8 |
| BF.24 | 22B (Omicron) | 8 |
| BE.1.1.2 | 22B (Omicron) | 8 |
| XE | recombinant | 8 |
| B.39 | 19A | 7 |
| B.1.1.317 | 20B | 7 |
| B.1.177.77 | 20E (EU1) | 7 |
| AY.45 | 21J (Delta) | 7 |
| AY.111 | 21J (Delta) | 7 |
| AY.113 | 21J (Delta) | 7 |
| BA.4.3 | 22A (Omicron) | 7 |
| BF.26 | 22B (Omicron) | 7 |
| BA.5.1.20 | 22B (Omicron) | 7 |
| B.40 | 19A | 6 |
| B.1.1.420 | 20B | 6 |
| AY.58 | 21I (Delta) | 6 |
| AY.65 | 21I (Delta) | 6 |
| AY.43.6 | 21J (Delta) | 6 |
| AY.39 | 21J (Delta) | 6 |
| AY.4.2.1 | 21J (Delta) | 6 |
| BA.2.3.2 | 21L (Omicron) | 6 |
| BA.2.14 | 21L (Omicron) | 6 |
| BA.4.2 | 22A (Omicron) | 6 |
| BA.5.2.33 | 22B (Omicron) | 6 |
| BE.1.1.1 | 22B (Omicron) | 6 |
| BA.5.2.30 | 22B (Omicron) | 6 |
| BF.11 | 22B (Omicron) | 6 |
| BF.18 | 22B (Omicron) | 6 |
| BA.5.2.6 | 22B (Omicron) | 6 |
| XD | recombinant | 6 |

| Pangolin lineage | Nextstrain (WHO) clades | Frequency |
|---|---|---|
| A.23.1 | 19B | 5 |
| B.1.258.3 | 20A | 5 |
| B.1.1.198 | 20B | 5 |
| B.1.1.44 | 20B | 5 |
| C.36.3 | 20D | 5 |
| AY.62 | 21I (Delta) | 5 |
| AY.39.1 | 21J (Delta) | 5 |
| AY.7.1 | 21J (Delta) | 5 |
| AY.4.9 | 21J (Delta) | 5 |
| BA.2.7 | 21L (Omicron) | 5 |
| BA.2.5 | 21L (Omicron) | 5 |
| B.1.1.529 | 21L (Omicron) | 5 |
| BA.5.1.4 | 22B (Omicron) | 5 |
| BA.5.1.21 | 22B (Omicron) | 5 |
| BA.5.1.25 | 22B (Omicron) | 5 |
| BA.5.2.7 | 22B (Omicron) | 5 |
| XM | recombinant | 5 |
| B.1.212 | 20A | 4 |
| B.1.398 | 20A | 4 |
| B.1.1.232 | 20B | 4 |
| B.1.351.5 | 20H (Beta, V2) | 4 |
| AY.4.6 | 21J (Delta) | 4 |
| BA.1.1.2 | 21K (Omicron) | 4 |
| BA.1.10 | 21K (Omicron) | 4 |
| BA.1.1.16 | 21K (Omicron) | 4 |
| BA.2.2 | 21L (Omicron) | 4 |
| BA.2.69 | 21L (Omicron) | 4 |
| BA.2.40.1 | 21L (Omicron) | 4 |
| BA.2.44 | 21L (Omicron) | 4 |
| BA.4.7 | 22A (Omicron) | 4 |
| BF.9 | 22B (Omicron) | 4 |
| BA.5.1.9 | 22B (Omicron) | 4 |
| BF.13 | 22B (Omicron) | 4 |
| BA.5.2.22 | 22B (Omicron) | 4 |
| BF.7.9 | 22B (Omicron) | 4 |
| XAR | recombinant | 4 |
| XAP | recombinant | 4 |
| XAF | recombinant | 4 |
| B.3 | 19A | 3 |
| B.1.236 | 20A | 3 |
| B.1.609 | 20A | 3 |
| B.1.596 | 20A | 3 |
| B.1.36.29 | 20A | 3 |
| B.1.1.406 | 20B | 3 |
| B.1.1.519 | 20B | 3 |
| B.1.177.10 | 20E (EU1) | 3 |
| P.1.1 | 20J (Gamma, V3) | 3 |

| Pangolin lineage | Nextstrain (WHO) clades | Frequency |
|---|---|---|
| AY.68 | 21I (Delta) | 3 |
| AY.44 | 21J (Delta) | 3 |
| AY.25.1 | 21J (Delta) | 3 |
| AY.122.1 | 21J (Delta) | 3 |
| AY.121.1 | 21J (Delta) | 3 |
| AY.127.1 | 21J (Delta) | 3 |
| BA.1.1.13 | 21K (Omicron) | 3 |
| BA.2.47 | 21L (Omicron) | 3 |
| BA.2.51 | 21L (Omicron) | 3 |
| BA.2.26 | 21L (Omicron) | 3 |
| BA.2.3.20 | 21L (Omicron) | 3 |
| BA.5.2.19 | 22B (Omicron) | 3 |
| BA.5.1.7 | 22B (Omicron) | 3 |
| BA.5.2.16 | 22B (Omicron) | 3 |
| BA.5.1.1 | 22B (Omicron) | 3 |
| BA.5.1.31 | 22B (Omicron) | 3 |
| BA.5.2.26 | 22B (Omicron) | 3 |
| BF.14 | 22B (Omicron) | 3 |
| BT.2 | 22B (Omicron) | 3 |
| CR.1 | 22B (Omicron) | 3 |
| BF.21 | 22B (Omicron) | 3 |
| BF.7.4 | 22B (Omicron) | 3 |
| BA.2.75.2 | 22D (Omicron) | 3 |
| XN | recombinant | 3 |
| XAC | recombinant | 3 |
| B.6 | 19A | 2 |
| B.1.22 | 20A | 2 |
| B.1.260 | 20A | 2 |
| B.1.438 | 20A | 2 |
| B.1.222 | 20A | 2 |
| B.1.379 | 20A | 2 |
| B.1.1.101 | 20B | 2 |
| B.1.533 | 20B | 2 |
| B.1.1.39 | 20B | 2 |
| B.1.1.294 | 20B | 2 |
| B.1.1.277 | 20B | 2 |
| B.1.1.153 | 20B | 2 |
| B.1.1.99 | 20B | 2 |
| B.1.362 | 20C | 2 |
| B.1.177.87 | 20E (EU1) | 2 |
| B.1.177.44 | 20E (EU1) | 2 |
| B.1.177.60 | 20E (EU1) | 2 |
| B.1.429 | 21C (Epsilon) | 2 |
| BB.2 | 21H (Mu) | 2 |
| AY.26 | 21I (Delta) | 2 |
| AY.3 | 21J (Delta) | 2 |
| AY.120.2.1 | 21J (Delta) | 2 |

| Pangolin lineage | Nextstrain (WHO) clades | Frequency |
|---|---|---|
| AY.87 | 21J (Delta) | 2 |
| AY.4.15 | 21J (Delta) | 2 |
| AY.46.4 | 21J (Delta) | 2 |
| AY.94 | 21J (Delta) | 2 |
| AY.78 | 21J (Delta) | 2 |
| AY.124.1 | 21J (Delta) | 2 |
| AY.25 | 21J (Delta) | 2 |
| AY.122.2 | 21J (Delta) | 2 |
| AY.43.5 | 21J (Delta) | 2 |
| AY.33.2 | 21J (Delta) | 2 |
| BA.1.6 | 21K (Omicron) | 2 |
| BA.1.15.3 | 21K (Omicron) | 2 |
| BA.2.24 | 21L (Omicron) | 2 |
| BA.2.16 | 21L (Omicron) | 2 |
| BA.2.27 | 21L (Omicron) | 2 |
| BA.2.32 | 21L (Omicron) | 2 |
| BA.2.3.9 | 21L (Omicron) | 2 |
| BA.2.30 | 21L (Omicron) | 2 |
| BA.2.11 | 21L (Omicron) | 2 |
| BA.2.9.6 | 21L (Omicron) | 2 |
| BA.2.13.1 | 21L (Omicron) | 2 |
| BA.2.10.3 | 21L (Omicron) | 2 |
| BS.1 | 21L (Omicron) | 2 |
| BA.4.1.4 | 22A (Omicron) | 2 |
| BA.4.1.8 | 22A (Omicron) | 2 |
| BA.5.2.4 | 22B (Omicron) | 2 |
| BF.25 | 22B (Omicron) | 2 |
| BA.5.2.28 | 22B (Omicron) | 2 |
| BF.3 | 22B (Omicron) | 2 |
| DE.1 | 22B (Omicron) | 2 |
| BA.5.2.8 | 22B (Omicron) | 2 |
| CC.1 | 22B (Omicron) | 2 |
| BF.7.6 | 22B (Omicron) | 2 |
| BA.2.75.3 | 22D (Omicron) | 2 |
| BL.1 | 22D (Omicron) | 2 |
| BM.1.1.3 | 22D (Omicron) | 2 |
| BQ.1.5 | 22E (Omicron) | 2 |
| XAB | recombinant | 2 |
| A.5 | 19B | 1 |
| A.19 | 19B | 1 |
| B.1.459 | 20A | 1 |
| B.1.9 | 20A | 1 |
| B.1.36.35 | 20A | 1 |
| B.1.214 | 20A | 1 |
| B.1.600 | 20A | 1 |
| B.1.78 | 20A | 1 |
| B.1.36.16 | 20A | 1 |

| Pangolin lineage | Nextstrain (WHO) clades | Frequency |
|---|---|---|
| B.1.243 | 20A | 1 |
| B.1.258.17 | 20A | 1 |
| B.1.619 | 20A | 1 |
| B.1.1.378 | 20B | 1 |
| B.1.1.61 | 20B | 1 |
| B.1.1.329 | 20B | 1 |
| B.1.1.231 | 20B | 1 |
| B.1.1.306 | 20B | 1 |
| B.1.1.351 | 20B | 1 |
| B.1.428 | 20C | 1 |
| B.1.623 | 20C | 1 |
| B.1.433 | 20C | 1 |
| B.1.575 | 20C | 1 |
| B.1.177.75 | 20E (EU1) | 1 |
| B.1.177.52 | 20E (EU1) | 1 |
| B.1.177.15 | 20E (EU1) | 1 |
| B.1.177.86 | 20E (EU1) | 1 |
| B.1.2 | 20G | 1 |
| P.1.17 | 20J (Gamma, V3) | 1 |
| AY.14 | 21A (Delta) | 1 |
| B.1.427 | 21C (Epsilon) | 1 |
| B.1.526 | 21F (Iota) | 1 |
| C.37 | 21G (Lambda) | 1 |
| AY.55 | 21I (Delta) | 1 |
| AY.24 | 21I (Delta) | 1 |
| AY.41 | 21J (Delta) | 1 |
| AY.128 | 21J (Delta) | 1 |
| AY.5.1 | 21J (Delta) | 1 |
| AY.119.2 | 21J (Delta) | 1 |
| AY.99.2 | 21J (Delta) | 1 |
| AY.125.1 | 21J (Delta) | 1 |
| AY.43.2 | 21J (Delta) | 1 |
| AY.98.1.1 | 21J (Delta) | 1 |
| AY.4.8 | 21J (Delta) | 1 |
| AY.84 | 21J (Delta) | 1 |
| AY.43.3 | 21J (Delta) | 1 |
| AY.4.17 | 21J (Delta) | 1 |
| AY.43.4 | 21J (Delta) | 1 |
| AY.4.13 | 21J (Delta) | 1 |
| BA.1.7 | 21K (Omicron) | 1 |
| BA.1.1.18 | 21K (Omicron) | 1 |
| BA.1.1.7 | 21K (Omicron) | 1 |
| BA.1.15.2 | 21K (Omicron) | 1 |
| BA.1.24 | 21K (Omicron) | 1 |
| BA.1.1.4 | 21K (Omicron) | 1 |
| BA.2.62 | 21L (Omicron) | 1 |
| BA.2.9.2 | 21L (Omicron) | 1 |

| Pangolin lineage | Nextstrain (WHO) clades | Frequency |
|---|---|---|
| BA.2.31 | 21L (Omicron) | 1 |
| BA.2.8 | 21L (Omicron) | 1 |
| BA.2.3.11 | 21L (Omicron) | 1 |
| BA.2.3.6 | 21L (Omicron) | 1 |
| BA.2.21 | 21L (Omicron) | 1 |
| BA.2.48 | 21L (Omicron) | 1 |
| BA.2.60 | 21L (Omicron) | 1 |
| BA.2.50 | 21L (Omicron) | 1 |
| BA.2.54 | 21L (Omicron) | 1 |
| BA.2.53 | 21L (Omicron) | 1 |
| BA.2.3.10 | 21L (Omicron) | 1 |
| BA.2.35 | 21L (Omicron) | 1 |
| BA.2.64 | 21L (Omicron) | 1 |
| BA.2.78 | 21L (Omicron) | 1 |
| BA.2.3.7 | 21L (Omicron) | 1 |
| BA.2.72 | 21L (Omicron) | 1 |
| BA.2.38.3 | 21L (Omicron) | 1 |
| BA.2.76 | 21L (Omicron) | 1 |
| BA.4.6.1 | 22A (Omicron) | 1 |
| BA.4.1.9 | 22A (Omicron) | 1 |
| BA.4.6.2 | 22A (Omicron) | 1 |
| BA.5.3.4 | 22B (Omicron) | 1 |
| BA.5.1.16 | 22B (Omicron) | 1 |
| BE.5 | 22B (Omicron) | 1 |
| BF.19 | 22B (Omicron) | 1 |
| BA.5.5.2 | 22B (Omicron) | 1 |
| BF.12 | 22B (Omicron) | 1 |
| BA.5.1.12 | 22B (Omicron) | 1 |
| BE.1.4 | 22B (Omicron) | 1 |
| BF.11.1 | 22B (Omicron) | 1 |
| BF.8 | 22B (Omicron) | 1 |
| BA.5.2.24 | 22B (Omicron) | 1 |
| BA.5.6.1 | 22B (Omicron) | 1 |
| BA.5.2.27 | 22B (Omicron) | 1 |
| BE.1.4.2 | 22B (Omicron) | 1 |
| BF.7.10 | 22B (Omicron) | 1 |
| BG.5 | 22C (Omicron) | 1 |
| BG.4 | 22C (Omicron) | 1 |
| BA.2.75.5 | 22D (Omicron) | 1 |
| BA.2.75.10 | 22D (Omicron) | 1 |
| BA.2.75.1 | 22D (Omicron) | 1 |
| BN.1.3 | 22D (Omicron) | 1 |
| BN.1 | 22D (Omicron) | 1 |
| BM.4.1.1 | 22D (Omicron) | 1 |
| BM.1.1 | 22D (Omicron) | 1 |
| BA.2.75.6 | 22D (Omicron) | 1 |
| BN.3 | 22D (Omicron) | 1 |

| Pangolin lineage | Nextstrain (WHO) clades | Frequency |
|---|---|---|
| BQ.1.16 | 22E (Omicron) | 1 |
| BQ.1.14 | 22E (Omicron) | 1 |
| BQ.1.10.1 | 22E (Omicron) | 1 |
| BQ.1.10 | 22E (Omicron) | 1 |
| BQ.1.15 | 22E (Omicron) | 1 |
| XS | recombinant | 1 |
| XV | recombinant | 1 |
| XAU | recombinant | 1 |
| XT | recombinant | 1 |
| XAH | recombinant | 1 |
| XAV | recombinant | 1 |
| XAK | recombinant | 1 |
| XBB.3 | recombinant | 1 |

**Supplementary Table S2. List of the nucleotide mutations by decreasing order of frequency among the 61,397 genomes**

| Nucleotide mutation | Total number of genomes harboring the nucleotide mutation |
|---|---|
| A23403G | 61267 |
| C14408U | 61234 |
| C3037U | 61188 |
| C241U | 53067 |
| C10029U | 43731 |
| C22995A | 40459 |
| G28881A | 28425 |
| G28883C | 28424 |
| G28882A | 28421 |
| C23604A | 27392 |
| G22992A | 26272 |
| U22917G | 25976 |
| A23063U | 25449 |
| A28271- | 25190 |
| C23525U | 23686 |
| G23948U | 23578 |
| C25584U | 23548 |
| U24469A | 23537 |
| A28271U | 23537 |
| C28311U | 23530 |
| U23599G | 23529 |
| 28362-28370 | 23522 |
| A24424U | 23509 |
| C26270U | 23506 |
| C10449A | 23505 |
| G26709A | 23377 |
| G22578A | 23140 |
| A18163G | 22804 |
| C22686U | 22465 |
| U22679C | 22402 |
| C22674U | 22236 |
| C23854A | 22220 |
| C27807U | 22121 |
| G29402U | 21571 |
| U26767C | 21437 |
| C23604G | 21404 |
| A23055G | 21367 |
| G15451A | 21335 |
| C25469U | 21313 |
| U23075C | 21290 |
| G28881U | 21250 |
| 28248-28253 | 21230 |
| C21618G | 21227 |
| G29742U | 21132 |
| A28461G | 21101 |
| A23013C | 20947 |
| C25000U | 20940 |
| C16466U | 20932 |
| 22029-22034 | 20671 |
| G210U | 20534 |
| G24410A | 20441 |
| C27752U | 20269 |
| C8986U | 20190 |
| G9053U | 20176 |
| G4181U | 20173 |
| A11201G | 20172 |
| C27874U | 20150 |
| C6402U | 20130 |
| A11332G | 20128 |
| C19220U | 20113 |
| U27638C | 19955 |
| 11288-11296 | 19901 |
| G21987A | 19755 |
| G28916U | 19570 |
| C7124U | 19564 |
| 21765-21770 | 18441 |
| C26577G | 18047 |
| G22813U | 17790 |
| A27259C | 17590 |
| C26060U | 15619 |
| C12880U | 15608 |
| C4321U | 15492 |
| C17410U | 15478 |
| C15714U | 15466 |
| C21618U | 15450 |
| 21633-21641 | 15421 |
| G4184A | 15407 |
| G10447A | 15401 |
| U22200G | 15387 |
| C2790U | 15371 |
| A20055G | 15364 |
| U670G | 15357 |
| C9534U | 15300 |
| A22786C | 15227 |
| A22688G | 15147 |

| Nucleotide mutation | Total number of genomes harboring the nucleotide mutation |
|---|---|
| C19955U | 15141 |
| G22775A | 15109 |
| A23040G | 15063 |
| C10198U | 15008 |
| A29510C | 14618 |
| C21846U | 13157 |
| C9344U | 12978 |
| A9424G | 12509 |
| C26858U | 9790 |
| U27384C | 9756 |
| G27382C | 9533 |
| A27383U | 9530 |
| G25563U | 9381 |
| C15952A | 9029 |
| A28299U | 8991 |
| C9866U | 8685 |
| C21762U | 8220 |
| C15240U | 8205 |
| A11537G | 8100 |
| C24503U | 8089 |
| G8393A | 8086 |
| 11285-11293 | 8083 |
| U13195C | 8079 |
| C24130A | 8074 |
| U5386G | 8069 |
| C23202A | 7972 |
| A2832G | 7943 |
| 21987-21995 | 7913 |
| A26530G | 7681 |
| 6513-6515 | 7530 |
| U22673C | 7147 |
| C18744U | 7092 |
| G23048A | 6881 |
| C25710U | 6764 |
| C18877U | 6616 |
| C26735U | 6581 |
| U22882G | 6563 |
| G5629U | 6528 |
| G9526U | 6506 |
| C11497U | 6504 |
| G13993U | 6503 |
| G15766U | 6503 |
| U26876C | 6494 |
| 22194-22196 | 6464 |
| C4543U | 6426 |
| G29399A | 6379 |
| G12160A | 6328 |
| G17019U | 6301 |
| A16889G | 6194 |
| C22792U | 6089 |
| G28975C | 6055 |
| U23018G | 5878 |
| G26529A | 5351 |
| C27889U | 5214 |
| G22599A | 4016 |
| 21992-21994 | 3991 |
| C14676U | 3947 |
| C27972U | 3932 |
| G28048U | 3932 |
| C5986U | 3924 |
| C913U | 3915 |
| C23709U | 3908 |
| C3267U | 3901 |
| C15279U | 3895 |
| U28282A | 3892 |
| A28281U | 3890 |
| G28280C | 3888 |
| G24914C | 3884 |
| C23271A | 3881 |
| U24506G | 3880 |
| C5388A | 3878 |
| U16176C | 3870 |
| C28977U | 3861 |
| A28111G | 3852 |
| U6954C | 3763 |
| G29734U | 3497 |
| 22204:GAGCCAGAA | 3200 |
| G28086U | 3048 |
| U27438C | 2714 |
| C29666U | 2707 |
| G526U | 2553 |
| G5437U | 2528 |
| G2518U | 2479 |
| C22314U | 2461 |
| C7768U | 2447 |

| Nucleotide mutation | Total number of genomes harboring the nucleotide mutation |
|---|---|
| A28330G | 2361 |
| C1627U | 2310 |
| C22227U | 2167 |
| C19524U | 2056 |
| G1048U | 1966 |
| G22898A | 1744 |
| C7851U | 1729 |
| C2470U | 1627 |
| C24130U | 1601 |
| G5924A | 1567 |
| A16064G | 1562 |
| C15237U | 1544 |
| G25855U | 1488 |
| C12073U | 1438 |
| C2416U | 1423 |
| C25872U | 1401 |
| C1059U | 1334 |
| C25624U | 1321 |
| A28095U | 1311 |
| C9891U | 1283 |
| C5730U | 1275 |
| C21575U | 1259 |
| A27038G | 1237 |
| C11514U | 1182 |
| G2258A | 1181 |
| C5622U | 1179 |
| C5184U | 1174 |
| G8371U | 1168 |
| G12310A | 1133 |
| U11418C | 1131 |
| C1758U | 1125 |
| C9924U | 1097 |
| G29778U | 1095 |
| G17488A | 1076 |
| G204U | 1072 |
| 29734-29759 | 1047 |
| C23683U | 1028 |
| G29557U | 1006 |
| U2746C | 999 |
| G28083U | 998 |
| A27745G | 953 |
| C22993U | 949 |
| U15096C | 942 |
| G8990U | 939 |
| 686-694 | 933 |
| G11083U | 926 |
| G25489U | 884 |
| C6286U | 862 |
| C23718U | 862 |
| C16616A | 836 |
| C6730U | 820 |
| C28253U | 812 |
| G21255C | 802 |
| U445C | 800 |
| G29645U | 783 |
| C28887U | 779 |
| U17040C | 777 |
| C26801G | 777 |
| G15906U | 773 |
| C23664U | 773 |
| A5584G | 772 |
| C28932U | 770 |
| C829U | 764 |
| G28878C | 759 |
| C27527U | 758 |
| A28877U | 758 |
| C15324U | 731 |
| C745U | 729 |
| G23593C | 715 |
| C26681U | 713 |
| G25471U | 712 |
| G20937U | 703 |
| C18086U | 699 |
| C27513U | 692 |
| C27012U | 668 |
| G1729A | 666 |
| G14030A | 662 |
| C13944U | 651 |
| A26786G | 645 |
| A10323G | 644 |
| G11851U | 642 |
| C13019U | 616 |
| G25785U | 605 |
| G18905A | 587 |
| G18255U | 574 |

| Nucleotide mutation | Total number of genomes harboring the nucleotide mutation |
|---|---|
| G29527U | 571 |
| G23012A | 564 |
| C1931A | 559 |
| C16466A | 553 |
| C21614U | 550 |
| C11020U | 548 |
| 28913-28915 | 547 |
| U25694C | 545 |
| A29700G | 543 |
| C25904U | 533 |
| A17615G | 532 |
| G29779U | 532 |
| 509-523 | 530 |
| C6196U | 529 |
| U21752C | 526 |
| C28724U | 519 |
| C10582U | 518 |
| C21855U | 516 |
| G5230U | 509 |
| C15738U | 509 |
| C9857U | 503 |
| U6979G | 502 |
| G28371U | 499 |
| C16887U | 477 |
| C29253U | 476 |
| G21800U | 474 |
| C29503U | 474 |
| C10525U | 471 |
| C25416U | 466 |
| G27014U | 465 |
| G28361U | 465 |
| C22498U | 460 |
| C2110U | 455 |
| C25824A | 453 |
| G26109A | 450 |
| C186U | 449 |
| C222U | 449 |
| G24928U | 448 |
| G9441U | 447 |
| C14120U | 444 |
| C28830A | 444 |
| G22104U | 441 |
| C27389U | 440 |
| G28001U | 437 |
| U17574C | 435 |
| C10279U | 431 |
| G28681U | 429 |
| G27516U | 428 |
| 28254 | 424 |
| C503U | 422 |
| C11750U | 422 |
| C25614U | 418 |
| G27788U | 415 |
| G22599C | 412 |
| C3096U | 407 |
| G16935A | 406 |
| C1191U | 400 |
| C6706U | 399 |
| C1912U | 398 |
| 25701:CCC | 394 |
| G17058U | 393 |
| C7564U | 386 |
| C4048U | 380 |
| U7106C | 378 |
| G487A | 377 |
| G6367U | 377 |
| U9283C | 377 |
| G26062U | 377 |
| U1877C | 376 |
| A21137G | 376 |
| A11782G | 373 |
| C25413U | 373 |
| G4960U | 364 |
| C5055U | 363 |
| U15521A | 362 |
| C10369U | 359 |
| G27261U | 358 |
| G174U | 355 |
| G29701A | 355 |
| G27632U | 354 |
| G25471C | 353 |
| G27906A | 352 |
| 510-518 | 350 |
| U21995C | 348 |
| A10646G | 347 |

| Nucleotide mutation | Total number of genomes harboring the nucleotide mutation |
|---|---|
| C6070U | 346 |
| C7303U | 346 |
| U21570G | 345 |
| C27804U | 345 |
| G23593U | 344 |
| G20060U | 340 |
| 515-520 | 339 |
| C4965U | 338 |
| G10688U | 337 |
| C29614U | 335 |
| C12784U | 334 |
| G19872U | 333 |
| C5392U | 331 |
| C26456U | 331 |
| G26720U | 331 |
| C29171U | 330 |
| A11595G | 328 |
| C3099U | 327 |
| G3764A | 327 |
| A20268G | 327 |
| A2692U | 326 |
| 22283-22291 | 326 |
| C9967U | 324 |
| C10301A | 324 |
| A21801C | 324 |
| A22206G | 324 |
| G24794U | 322 |
| A6517U | 321 |
| C9246U | 319 |
| C27143U | 318 |
| C14230A | 315 |
| G25352U | 313 |
| C5672U | 311 |
| C16575U | 310 |
| G29543U | 310 |
| G3638U | 309 |
| G24812U | 309 |
| G10642U | 307 |
| U11049C | 305 |
| C11747U | 303 |
| C22281U | 303 |
| 22205:GAGCCAGAA | 302 |
| C21595U | 301 |
| U24079C | 301 |
| A12755G | 300 |
| G23401U | 300 |
| G28884C | 300 |
| C25517U | 299 |
| G29628A | 299 |
| 518-520 | 298 |
| A5250G | 298 |
| C19983U | 296 |
| C19245U | 293 |
| A29403U | 292 |
| C6633U | 290 |
| C27549U | 290 |
| C13721U | 289 |
| G17259U | 289 |
| G24834A | 288 |
| C18377U | 286 |
| C5944U | 285 |
| C26029A | 285 |
| C28312U | 284 |
| U606C | 283 |
| G1820A | 283 |
| G9332A | 283 |
| C8991U | 282 |
| G7936U | 279 |
| C25810U | 279 |
| C5079U | 278 |
| C6040U | 278 |
| 27673-27696 | 278 |
| G27933A | 277 |
| G26526U | 276 |
| G1738U | 275 |
| G24872U | 274 |
| G28122U | 273 |
| U2954C | 269 |
| G29081U | 268 |
| G3764U | 267 |
| C26013U | 267 |
| C29311U | 266 |
| G29690U | 263 |
| C25513U | 261 |
| U23489C | 260 |

| Nucleotide mutation | Total number of genomes harboring the nucleotide mutation |
|---|---|
| C23191U | 258 |
| C678U | 257 |
| C5170U | 257 |
| C17285U | 257 |
| C3602U | 255 |
| G25996U | 255 |
| C26151U | 255 |
| C23673U | 249 |
| U8053C | 248 |
| U28140G | 248 |
| C27612U | 247 |
| G8150A | 245 |
| C25521U | 245 |
| G7925A | 244 |
| C19875U | 244 |
| G4006U | 243 |
| C22311U | 243 |
| G11132U | 242 |
| G11417U | 242 |
| G17427U | 240 |
| C8208U | 239 |
| C14724U | 239 |
| G22801U | 236 |
| G23126U | 236 |
| G13459A | 235 |
| C13957U | 235 |
| U26609C | 234 |
| G25311U | 232 |
| C27739U | 232 |
| A21647G | 231 |
| C23557U | 231 |
| C21742U | 230 |
| C21110U | 229 |
| C4276U | 227 |
| C8668U | 227 |
| C12789U | 226 |
| C25549U | 226 |
| G16853U | 224 |
| U26870C | 224 |
| C13297U | 222 |
| C6317U | 219 |
| C934U | 217 |
| C25844U | 217 |
| C26895U | 217 |
| C583U | 216 |
| C28651U | 216 |
| C28854U | 216 |
| G10396U | 215 |
| G25599U | 215 |
| A29110U | 215 |
| C3241U | 214 |
| C14599U | 212 |
| C11674U | 210 |
| G25690U | 210 |
| G19825U | 209 |
| C29095U | 207 |
| C22879U | 206 |
| C25708U | 206 |
| C27942U | 206 |
| C29686U | 206 |
| A20262G | 205 |
| G13921U | 204 |
| A14457G | 204 |
| C14925U | 204 |
| G18816U | 204 |
| C25626U | 204 |
| C27434U | 204 |
| C22458U | 203 |
| C24707A | 203 |
| C25886U | 203 |
| A6342G | 202 |
| U5071C | 201 |
| C16289U | 201 |
| G28079U | 201 |
| C7528U | 200 |
| C27944U | 200 |
| C1267U | 198 |
| C6027U | 198 |
| C12970U | 198 |
| C23188U | 197 |
| C6781U | 196 |
| C29769U | 196 |
| U28149C | 195 |
| C13255U | 194 |
| C13620U | 194 |

| Nucleotide mutation | Total number of genomes harboring the nucleotide mutation |
|---|---|
| G15372U | 194 |
| C19602U | 194 |
| A23767G | 194 |
| A29767U | 194 |
| C203U | 193 |
| U16371C | 193 |
| C28833U | 193 |
| G26995U | 192 |
| C307U | 191 |
| C2509U | 191 |
| A12162G | 191 |
| C24138U | 191 |
| C26111U | 191 |
| C13860U | 190 |
| G22225U | 190 |
| G25135U | 190 |
| C7056U | 189 |
| C25792U | 189 |
| C8140U | 188 |
| G11365U | 188 |
| G22132U | 188 |
| G28487A | 188 |
| G21641U | 187 |
| G23663U | 187 |
| G29511U | 187 |
| A4964G | 186 |
| C16726U | 186 |
| C21191U | 185 |
| C29750U | 185 |
| C10789U | 183 |
| U18018C | 183 |
| A4353G | 182 |
| A17207G | 182 |
| G15910U | 181 |
| A20713C | 181 |
| G29702A | 181 |
| C29738U | 181 |
| C2485U | 180 |
| C4893U | 180 |
| U24760C | 180 |
| U25473C | 180 |
| G24620U | 179 |
| G25621U | 179 |
| A28254C | 178 |
| C683U | 177 |
| A17337G | 177 |
| U22917A | 177 |
| A3291G | 176 |
| C8092U | 176 |
| C17676U | 176 |
| C26645U | 176 |
| C27532U | 176 |
| C28603U | 176 |
| C8251U | 175 |
| U16056C | 175 |
| G5504A | 174 |
| C8389U | 174 |
| C21721U | 174 |
| U27534C | 174 |
| C5284U | 173 |
| A22812C | 172 |
| C27577U | 172 |
| C28948U | 172 |
| G28378U | 171 |
| G21974U | 170 |
| C29679U | 170 |
| G7405U | 169 |
| G4904U | 168 |
| C6285U | 168 |
| C10632U | 168 |
| G25088U | 168 |
| C29445U | 168 |
| C1952A | 167 |
| A25505G | 167 |
| C4331U | 166 |
| A8052G | 166 |
| G27877U | 166 |
| A5648C | 165 |
| G20356A | 165 |
| G21088U | 165 |
| C23707U | 165 |
| C28432U | 165 |
| G7042U | 164 |
| G17411U | 164 |
| G28077A | 164 |

| Nucleotide mutation | Total number of genomes harboring the nucleotide mutation |
|---|---|
| C28269A | 164 |
| C29284U | 164 |
| C29520U | 164 |
| U12946C | 163 |
| C16111U | 163 |
| G1463U | 162 |
| C16293U | 162 |
| C20719U | 162 |
| C25587U | 162 |
| C28961U | 162 |
| C7420U | 161 |
| G1043U | 160 |
| C6539U | 160 |
| C17703U | 160 |
| C18348U | 160 |
| G19999U | 160 |
| C25613U | 160 |
| C28869U | 160 |
| C3738U | 159 |
| U8914C | 159 |
| U15009C | 158 |
| C28087U | 158 |
| C17470U | 157 |
| C6941U | 156 |
| G28077U | 156 |
| C29632U | 156 |
| C2453U | 155 |
| U20310C | 155 |
| G28851U | 155 |
| G22021U | 154 |
| C25096U | 154 |
| G25906C | 154 |
| C335U | 153 |
| G5210A | 153 |
| G28857U | 153 |
| U27125C | 152 |
| C18176U | 151 |
| C20762U | 151 |
| A23148G | 151 |
| G28198U | 151 |
| C1170U | 150 |
| C1684U | 150 |
| A23535G | 150 |
| G29706U | 150 |
| G5950U | 149 |
| G28178U | 149 |
| U7984C | 148 |
| C10138U | 148 |
| C15960U | 148 |
| U21765C | 148 |
| G25483A | 147 |
| A26003U | 147 |
| G25563C | 146 |
| G25740U | 146 |
| C230U | 145 |
| C1625U | 144 |
| G3871U | 144 |
| C15720U | 143 |
| C17403U | 143 |
| C18171U | 142 |
| G21305A | 142 |
| G22081U | 142 |
| U24224C | 142 |
| C28657U | 142 |
| U733C | 141 |
| C25355U | 141 |
| A3082G | 140 |
| C5907U | 140 |
| A20553G | 140 |
| C21658U | 140 |
| G1181U | 139 |
| C13665U | 139 |
| C27294U | 139 |
| C6471U | 138 |
| C12823U | 138 |
| U25975C | 138 |
| C10507U | 137 |
| C11455U | 137 |
| C24642U | 137 |
| C27059U | 137 |
| C29659U | 137 |
| C26801U | 136 |
| C2037U | 135 |
| C2706U | 135 |
| C12068U | 135 |

| Nucleotide mutation | Total number of genomes harboring the nucleotide mutation |
|---|---|
| C17104U | 135 |
| C22088U | 135 |
| C25731U | 135 |
| C9565U | 134 |
| C7162U | 133 |
| C9430U | 133 |
| C18555U | 133 |
| C21034U | 133 |
| C21638U | 133 |
| A24614G | 133 |
| G14500U | 132 |
| C23230U | 132 |
| U10135C | 131 |
| G11335U | 131 |
| C12778U | 131 |
| C16323U | 131 |
| C19374U | 131 |
| G25437U | 131 |
| G25538U | 131 |
| C3828U | 129 |
| C8917U | 129 |
| C27393U | 129 |
| A27620U | 129 |
| C28093U | 129 |
| C28462U | 129 |
| G28655U | 129 |
| C4113U | 128 |
| C10977U | 128 |
| C10039U | 127 |
| A14125G | 127 |
| G22468U | 127 |
| C22480U | 127 |
| C14407U | 126 |
| C26408U | 126 |
| U27071C | 126 |
| A29301G | 126 |
| C15656U | 125 |
| C6651U | 124 |
| C12025U | 124 |
| C12915U | 124 |
| C28435U | 124 |
| A1643U | 123 |
| C4320U | 123 |
| C5183U | 123 |
| C11824U | 123 |
| G28167A | 123 |
| U28251C | 123 |
| C29642U | 123 |
| C6310U | 122 |
| C9661U | 122 |
| G10870U | 122 |
| G25726U | 122 |
| G29179U | 122 |
| C12786U | 121 |
| U14257C | 121 |
| C21304A | 121 |
| C28253G | 121 |
| 28256:UC | 121 |
| G28878A | 121 |
| G29711U | 121 |
| G625U | 120 |
| C3874U | 120 |
| G5194A | 120 |
| C27643U | 119 |
| A11781G | 118 |
| C13458U | 118 |
| C21219U | 118 |
| C23248U | 118 |
| U23560C | 118 |
| C313U | 117 |
| C1528U | 117 |
| C24748U | 117 |
| C25889U | 117 |
| C4084U | 116 |
| C20946U | 116 |
| G21786U | 116 |
| C22597U | 116 |
| A27218G | 116 |
| C2749U | 115 |
| C2910U | 115 |
| G3892A | 115 |
| C8782U | 115 |
| G14743A | 115 |
| G3606U | 114 |
| C3787U | 114 |

| Nucleotide mutation | Total number of genomes harboring the nucleotide mutation |
|---|---|
| A6613G | 114 |
| U8603C | 114 |
| A17039G | 114 |
| C25207U | 114 |
| C28531U | 114 |
| C6380U | 113 |
| C7224U | 113 |
| U26149C | 113 |
| C27575U | 113 |
| C5884U | 112 |
| A6319G | 112 |
| C7764U | 112 |
| C9223U | 112 |
| C20703U | 112 |
| C26256U | 112 |
| C3177U | 111 |
| G11330A | 111 |
| C12119U | 111 |
| C1498U | 110 |
| C17304U | 110 |
| C19164U | 110 |
| C19269U | 110 |
| U22942A | 110 |
| G24390C | 110 |
| C25658U | 110 |
| C28512G | 110 |
| A1807G | 109 |
| C5512U | 109 |
| C16193U | 109 |
| C21621A | 109 |
| C23647U | 109 |
| G27505U | 109 |
| A28699G | 109 |
| G2030U | 108 |
| A3859G | 108 |
| C4586U | 108 |
| C19862U | 108 |
| G22103C | 108 |
| G2095A | 107 |
| C2232U | 107 |
| G4913A | 107 |
| C4940U | 107 |
| G5425U | 107 |
| G6513A | 107 |
| G12769A | 107 |
| G16377U | 107 |
| C18508U | 107 |
| G25340A | 107 |
| G25534U | 107 |
| G26199U | 107 |
| U29223C | 107 |
| G29747U | 107 |
| G2659A | 106 |
| C7869U | 106 |
| U8593C | 106 |
| C20428U | 106 |
| A20724G | 106 |
| G22205U | 106 |
| C24865U | 106 |
| 28278-28280 | 106 |
| C29375A | 106 |
| G571A | 105 |
| G2198A | 105 |
| C6401U | 105 |
| U9454C | 105 |
| G11114U | 105 |
| G22894A | 105 |
| G26828U | 105 |
| C28308G | 105 |
| U2698C | 104 |
| A10286G | 104 |
| G11410C | 104 |
| G19563A | 104 |
| G20679U | 104 |
| A24110C | 104 |
| C5812U | 103 |
| C12049U | 103 |
| C22432U | 103 |
| A25439C | 103 |
| C25700U | 103 |
| 27205-27207 | 103 |
| U28297C | 103 |
| C5822U | 102 |
| C19011U | 102 |
| C20628U | 102 |

| Nucleotide mutation | Total number of genomes harboring the nucleotide mutation |
|---|---|
| A24389C | 102 |
| G26428U | 102 |
| C29774U | 102 |
| G4201U | 101 |
| C8407U | 101 |
| C9142U | 101 |
| C12781U | 101 |
| C14757U | 101 |
| G17278U | 101 |
| C18115U | 101 |
| G18210U | 101 |
| G21824C | 101 |
| 28262:AACA | 101 |
| A28402G | 101 |
| C29708U | 101 |
| G29737U | 101 |
| G17686U | 100 |
| C18129U | 100 |
| G20006U | 100 |
| C24378U | 100 |
| G29751U | 100 |
| C2695U | 99 |
| G27816A | 99 |
| U28144C | 99 |
| A12417G | 98 |
| A17799G | 98 |
| U18465C | 98 |
| C19017U | 98 |
| C19390U | 98 |
| C20104U | 98 |
| G26167U | 98 |
| G27390U | 98 |
| C28677U | 98 |
| C29409U | 98 |
| G29751A | 98 |
| G29781U | 98 |
| C6128U | 97 |
| C7749U | 97 |
| G12911A | 97 |
| C13536U | 97 |
| C17012U | 97 |
| G21520U | 97 |
| C22326U | 97 |
| C22747U | 97 |
| G29734C | 97 |
| 516-518 | 96 |
| G2210U | 96 |
| C2902U | 96 |
| A14107G | 96 |
| C14913U | 96 |
| G15327U | 96 |
| C19366U | 96 |
| G19549U | 96 |
| A21717G | 96 |
| G25218U | 96 |
| G29736U | 96 |
| G1006U | 95 |
| C2455U | 95 |
| C3743U | 95 |
| U4579A | 95 |
| C7164U | 95 |
| C13724U | 95 |
| G18186U | 95 |
| C23638U | 95 |
| C27630U | 95 |
| C27657U | 95 |
| U27696C | 95 |
| C337U | 94 |
| C823U | 94 |
| G1228A | 94 |
| C1281U | 94 |
| C1348U | 94 |
| G4510U | 94 |
| C4582U | 94 |
| C6541U | 94 |
| C7113U | 94 |
| C8950U | 94 |
| C12115U | 94 |
| G14371U | 94 |
| U15684C | 94 |
| C21057U | 94 |
| C22858U | 94 |
| G24821U | 94 |
| G25552U | 94 |
| G27703U | 94 |

| Nucleotide mutation | Total number of genomes harboring the nucleotide mutation |
|---|---|
| C2189U | 93 |
| C6582U | 93 |
| C9559U | 93 |
| C9803U | 93 |
| A18468G | 93 |
| C22323U | 93 |
| U23042C | 93 |
| C23086U | 93 |
| G25445A | 93 |
| G29742A | 93 |
| U9508C | 92 |
| A21532G | 92 |
| C21727U | 92 |
| G22017U | 92 |
| C25578U | 92 |
| C29366U | 92 |
| C29635U | 92 |
| A4681G | 91 |
| G8137U | 91 |
| A16044U | 91 |
| C16329U | 91 |
| C21789U | 91 |
| A22012U | 91 |
| U26075C | 91 |
| U26076A | 91 |
| G28895U | 91 |
| A29382G | 91 |
| C4752U | 90 |
| C26305U | 90 |
| C835U | 89 |
| A1098G | 89 |
| G1957U | 89 |
| C2113U | 89 |
| C5826U | 89 |
| C5850U | 89 |
| C9693U | 89 |
| G14209U | 89 |
| C19170U | 89 |
| C21622U | 89 |
| C24138A | 89 |
| G24199U | 89 |
| C106U | 88 |
| C1288U | 88 |
| C1513U | 88 |
| C2445U | 88 |
| C8326U | 88 |
| A9241G | 88 |
| C9451U | 88 |
| C11956U | 88 |
| C12008U | 88 |
| G15543U | 88 |
| A28384U | 88 |
| C832U | 87 |
| C1420U | 87 |
| C6026U | 87 |
| C6936U | 87 |
| U9376C | 87 |
| C14838U | 87 |
| A21222U | 87 |
| C22120U | 87 |
| C24210U | 87 |
| C27707U | 87 |
| G28727U | 87 |
| C1616U | 86 |
| G2600U | 86 |
| G3483U | 86 |
| C21597U | 86 |
| C22033A | 86 |
| C24904U | 86 |
| G25687U | 86 |
| C27476U | 86 |
| C27945U | 86 |
| U29728G | 86 |
| G598A | 85 |
| G3620A | 85 |
| A6672G | 85 |
| 6686-6688 | 85 |
| G8189A | 85 |
| A9852G | 85 |
| C9866A | 85 |
| C11758U | 85 |
| C12459U | 85 |
| C20270U | 85 |
| C22388U | 85 |
| C24912U | 85 |

| Nucleotide mutation | Total number of genomes harboring the nucleotide mutation |
|---|---|
| C27005U | 85 |
| G27561U | 85 |
| C449U | 84 |
| C799U | 84 |
| G1274A | 84 |
| C1812U | 84 |
| C8047U | 84 |
| U8885C | 84 |
| C12855U | 84 |
| C1437U | 83 |
| C2334U | 83 |
| C6312U | 83 |
| A8530G | 83 |
| C11109U | 83 |
| C18788U | 83 |
| C21058U | 83 |
| C21811U | 83 |
| C25047U | 83 |
| U26035C | 83 |
| A1585C | 82 |
| C4540U | 82 |
| C5144U | 82 |
| G8179U | 82 |
| C10702U | 82 |
| C12747U | 82 |
| G19518U | 82 |
| 22196-22198 | 82 |
| U25711C | 82 |
| G27754U | 82 |
| C1594U | 81 |
| C4456U | 81 |
| G8102U | 81 |
| C11152U | 81 |
| G16647A | 81 |
| G19509A | 81 |
| C25728U | 81 |
| G28798A | 81 |
| C29272U | 81 |
| C29358U | 81 |
| C1426U | 80 |
| C3903U | 80 |
| C5365U | 80 |
| C10376U | 80 |
| U19839C | 80 |
| G25340U | 80 |
| G29392U | 80 |
| C412U | 79 |
| C999A | 79 |
| C3653U | 79 |
| U4690C | 79 |
| C9073U | 79 |
| A9419G | 79 |
| C12513U | 79 |
| G15372C | 79 |
| C17999U | 79 |
| C24370U | 79 |
| U25333C | 79 |
| C28344U | 79 |
| G29262U | 79 |
| C364U | 78 |
| A2941G | 78 |
| G3563A | 78 |
| C3876U | 78 |
| A5662G | 78 |
| G7925U | 78 |
| C10252U | 78 |
| A20003C | 78 |
| C20844U | 78 |
| C21077U | 78 |
| C25511U | 78 |
| C26549U | 78 |
| C28725U | 78 |
| G29688U | 78 |
| C1938U | 77 |
| C3811U | 77 |
| C4002U | 77 |
| C9286U | 77 |
| C10615U | 77 |
| C11620U | 77 |
| C14184U | 77 |
| G18756U | 77 |
| C20451U | 77 |
| 22198:UCCGGCAGA | 77 |
| C22642U | 77 |
| G22918U | 77 |

| Nucleotide mutation | Total number of genomes harboring the nucleotide mutation |
|---|---|
| G27706U | 77 |
| C29535U | 77 |
| C29627U | 77 |
| C346U | 76 |
| C635U | 76 |
| C1551U | 76 |
| C4927U | 76 |
| C5526U | 76 |
| U7767C | 76 |
| C11173U | 76 |
| C15579U | 76 |
| U18358C | 76 |
| A19667G | 76 |
| C20031U | 76 |
| C21707U | 76 |
| C22987U | 76 |
| C24083U | 76 |
| C27600U | 76 |
| C27741U | 76 |
| C29451U | 76 |
| G1444A | 75 |
| A10884G | 75 |
| C11752U | 75 |
| C12439U | 75 |
| G14497A | 75 |
| G20709U | 75 |
| G22346U | 75 |
| C25916U | 75 |
| C26753U | 75 |
| C28849U | 75 |
| C2675U | 74 |
| C3040U | 74 |
| C4777U | 74 |
| C13168U | 74 |
| G16827A | 74 |
| G18028U | 74 |
| C18189U | 74 |
| C18395U | 74 |
| G21795U | 74 |
| G22899U | 74 |
| C24023U | 74 |
| C25046U | 74 |
| G25455U | 74 |
| G25537U | 74 |
| G25654U | 74 |
| C25688U | 74 |
| U26442C | 74 |
| C27442U | 74 |
| G29766U | 74 |
| 29767-29776 | 74 |
| G2305A | 73 |
| G3875A | 73 |
| C5869U | 73 |
| C6255U | 73 |
| C11704U | 73 |
| C12473U | 73 |
| G15768U | 73 |
| C18486U | 73 |
| C18981U | 73 |
| C21304U | 73 |
| U21632C | 73 |
| U23317C | 73 |
| G24368U | 73 |
| G25445U | 73 |
| C25572U | 73 |
| G25720U | 73 |
| C27297U | 73 |
| G28209U | 73 |
| C29762U | 73 |
| C2306U | 72 |
| C5628U | 72 |
| G7393U | 72 |
| C7735U | 72 |
| C7765U | 72 |
| A8882G | 72 |
| U9875C | 72 |
| C10416U | 72 |
| U18402C | 72 |
| C22000U | 72 |
| C22916A | 72 |
| C26885U | 72 |
| G28541U | 72 |
| G29254U | 72 |
| 3104-3109 | 71 |
| A6497G | 71 |

| Nucleotide mutation | Total number of genomes harboring the nucleotide mutation |
|---|---|
| C10156U | 71 |
| G15780A | 71 |
| C16247U | 71 |
| G16622A | 71 |
| C17746U | 71 |
| C21627U | 71 |
| C24382U | 71 |
| C24421U | 71 |
| U25016C | 71 |
| C25276U | 71 |
| C27247U | 71 |
| G28985U | 71 |
| C29754U | 71 |
| C3393U | 70 |
| C3426U | 70 |
| G4162U | 70 |
| C5310U | 70 |
| C6720U | 70 |
| C6896U | 70 |
| A11822G | 70 |
| C16049U | 70 |
| C16308U | 70 |
| C20178U | 70 |
| G21296A | 70 |
| C22450U | 70 |
| C25006U | 70 |
| G25459U | 70 |
| C25611U | 70 |
| 28090-28095 | 70 |
| C28354U | 70 |
| C28744U | 70 |
| G219U | 69 |
| C1122U | 69 |
| C4763U | 69 |
| C5526A | 69 |
| C6843U | 69 |
| C8016U | 69 |
| C13225U | 69 |
| G14118U | 69 |
| C16376U | 69 |
| U17139A | 69 |
| G18105U | 69 |
| A18366G | 69 |
| G19684U | 69 |
| C19895U | 69 |
| G20208U | 69 |
| G20580U | 69 |
| A22320G | 69 |
| G27621A | 69 |
| G28027U | 69 |
| C29585U | 69 |
| C1387U | 68 |
| C2973U | 68 |
| C3176U | 68 |
| U4917C | 68 |
| A5995G | 68 |
| U9861C | 68 |
| C10440U | 68 |
| G18181A | 68 |
| C21952U | 68 |
| C22224U | 68 |
| C25721U | 68 |
| C29485U | 68 |
| C1404U | 67 |
| C4832U | 67 |
| C6573U | 67 |
| G6622A | 67 |
| C6990U | 67 |
| C10728U | 67 |
| G11521A | 67 |
| C12076U | 67 |
| A12410G | 67 |
| C13806U | 67 |
| C14805U | 67 |
| C18457U | 67 |
| G19962U | 67 |
| G23675A | 67 |
| U24892C | 67 |
| C29200U | 67 |
| C3768U | 66 |
| G4300U | 66 |
| C6388U | 66 |
| C6555U | 66 |
| C6968U | 66 |
| C7834U | 66 |

| Nucleotide mutation | Total number of genomes harboring the nucleotide mutation |
|---|---|
| C14178U | 66 |
| C16362U | 66 |
| C17550U | 66 |
| C18175U | 66 |
| C18252U | 66 |
| G20131A | 66 |
| C20759U | 66 |
| C20930U | 66 |
| 22031-22036 | 66 |
| C22311A | 66 |
| U22672C | 66 |
| C22879A | 66 |
| U24499C | 66 |
| C26882U | 66 |
| U27484C | 66 |
| 27792-27793 | 66 |
| G28903U | 66 |
| C29077U | 66 |
| G29140U | 66 |
| C1473U | 65 |
| C1593U | 65 |
| C3369U | 65 |
| A4333G | 65 |
| C4891U | 65 |
| C6543U | 65 |
| C13694U | 65 |
| C17004U | 65 |
| G17014U | 65 |
| C19185U | 65 |
| U21982C | 65 |
| C23202U | 65 |
| G23402A | 65 |
| C24442U | 65 |
| U27656G | 65 |
| G29449U | 65 |
| G29628U | 65 |
| C2197U | 64 |
| A2565G | 64 |
| A3252G | 64 |
| C7086U | 64 |
| C7165U | 64 |
| C8290U | 64 |
| C9319U | 64 |
| C24374U | 64 |
| C26606U | 64 |
| G27463U | 64 |
| C27509U | 64 |
| C27944G | 64 |
| A28521G | 64 |
| C28708U | 64 |
| U28726A | 64 |
| G587A | 63 |
| C3264U | 63 |
| U6235C | 63 |
| C8964U | 63 |
| A13756G | 63 |
| C25463U | 63 |
| G25855C | 63 |
| G26122A | 63 |
| C28453U | 63 |
| G29736C | 63 |
| C1441U | 62 |
| C8293U | 62 |
| G10097A | 62 |
| C10969U | 62 |
| C11200U | 62 |
| C11653U | 62 |
| C14220U | 62 |
| G14398U | 62 |
| G18148A | 62 |
| C18312A | 62 |
| C18570U | 62 |
| A19257G | 62 |
| U21527C | 62 |
| C22591U | 62 |
| C23613U | 62 |
| U25353C | 62 |
| C26054A | 62 |
| G26211U | 62 |
| G29777U | 62 |
| C2644U | 61 |
| C6701U | 61 |
| C7600U | 61 |
| C10277U | 61 |
| C11094U | 61 |

| Nucleotide mutation | Total number of genomes harboring the nucleotide mutation |
|---|---|
| C15540U | 61 |
| G17331U | 61 |
| C17634U | 61 |
| C17977U | 61 |
| G22335U | 61 |
| C23757U | 61 |
| G26153U | 61 |
| C27800A | 61 |
| C28498U | 61 |
| C28697U | 61 |
| C28775U | 61 |
| G28882U | 61 |
| G28899U | 61 |
| C29296U | 61 |
| G219A | 60 |
| C593U | 60 |
| G2398A | 60 |
| G4399U | 60 |
| A5703C | 60 |
| C7267U | 60 |
| C13342U | 60 |
| U20100C | 60 |
| G20292A | 60 |
| C20629U | 60 |
| U20911C | 60 |
| G21468U | 60 |
| C23731U | 60 |
| G24757U | 60 |
| G25906U | 60 |
| C26625U | 60 |
| U26972C | 60 |
| C27493U | 60 |
| C27494U | 60 |
| C28054U | 60 |
| C28310U | 60 |
| A29139U | 60 |
| C29541U | 60 |
| C1205G | 59 |
| G2782U | 59 |
| C3330U | 59 |
| C3817U | 59 |
| C5806U | 59 |
| C10834U | 59 |
| C11941U | 59 |
| G12131A | 59 |
| C15654U | 59 |
| G15672U | 59 |
| G16744A | 59 |
| C17010U | 59 |
| C17474U | 59 |
| C17518U | 59 |
| C17747U | 59 |
| C19586U | 59 |
| C22624U | 59 |
| G25606U | 59 |
| G26951C | 59 |
| C29364U | 59 |
| C29509U | 59 |
| C29733U | 59 |
| G29764A | 59 |
| C1514U | 58 |
| C3261U | 58 |
| U4237C | 58 |
| C4455U | 58 |
| C5239U | 58 |
| C5974U | 58 |
| C6578U | 58 |
| G8174A | 58 |
| C8299U | 58 |
| C10537U | 58 |
| C12525U | 58 |
| G17122U | 58 |
| C17766U | 58 |
| C18032U | 58 |
| C19884U | 58 |
| C20133U | 58 |
| C21306U | 58 |
| U25959C | 58 |
| C26028U | 58 |
| C26313U | 58 |
| C29555U | 58 |
| C2062U | 57 |
| A2647G | 57 |
| G3431U | 57 |
| G4207U | 57 |

| Nucleotide mutation | Total number of genomes harboring the nucleotide mutation |
|---|---|
| C8660U | 57 |
| C8947U | 57 |
| C10116U | 57 |
| C10336A | 57 |
| A17858G | 57 |
| C21646U | 57 |
| U24910C | 57 |
| C25350U | 57 |
| U25518C | 57 |
| C25714U | 57 |
| C27737U | 57 |
| C29167U | 57 |
| U29335C | 57 |
| U365A | 56 |
| A2521G | 56 |
| C2536U | 56 |
| A2786G | 56 |
| A3684G | 56 |
| C5339U | 56 |
| G5515U | 56 |
| U6148C | 56 |
| C7119U | 56 |
| A8031G | 56 |
| A13300U | 56 |
| C13335U | 56 |
| G14511C | 56 |
| C18657U | 56 |
| C18804U | 56 |
| U25783C | 56 |
| G27604U | 56 |
| C28606U | 56 |
| U29029A | 56 |
| C29370U | 56 |
| A29776U | 56 |
| U580C | 55 |
| G3965U | 55 |
| G7675U | 55 |
| C8344U | 55 |
| C9811U | 55 |
| C9943U | 55 |
| G12769U | 55 |
| G13812U | 55 |
| C14318U | 55 |
| C15810U | 55 |
| A15936G | 55 |
| C16428U | 55 |
| C17934U | 55 |
| C18029U | 55 |
| C20404U | 55 |
| C22295U | 55 |
| G23012C | 55 |
| G25644U | 55 |
| G27762U | 55 |
| C28057U | 55 |
| G29422U | 55 |
| A29500G | 55 |
| C228U | 54 |
| C900U | 54 |
| C2623U | 54 |
| C3784U | 54 |
| C9166U | 54 |
| G15598A | 54 |
| G17658U | 54 |
| C19151U | 54 |
| G19735U | 54 |
| C25553U | 54 |
| C25603U | 54 |
| A28129G | 54 |
| C28291U | 54 |
| G29260A | 54 |
| G2185U | 53 |
| G5023U | 53 |
| C5575U | 53 |
| C6336U | 53 |
| A6604G | 53 |
| C8078U | 53 |
| C9165U | 53 |
| G9832U | 53 |
| G12988U | 53 |
| C13274U | 53 |
| C15660U | 53 |
| C16260U | 53 |
| C18060U | 53 |
| G18079U | 53 |
| C18647U | 53 |

| Nucleotide mutation | Total number of genomes harboring the nucleotide mutation |
|---|---|
| C21781U | 53 |
| C22377U | 53 |
| C23635U | 53 |
| C25460U | 53 |
| C27002U | 53 |
| C27526U | 53 |
| C29149U | 53 |
| C601U | 52 |
| C1457U | 52 |
| U1885A | 52 |
| A1964G | 52 |
| C3511U | 52 |
| C8240U | 52 |
| C11650U | 52 |
| U13048C | 52 |
| C14649U | 52 |
| C16338U | 52 |
| A16482G | 52 |
| C17125U | 52 |
| A17739G | 52 |
| U18545C | 52 |
| C19854U | 52 |
| C20148U | 52 |
| U20407C | 52 |
| G21220U | 52 |
| G21624U | 52 |
| U23031C | 52 |
| G23120U | 52 |
| C26907U | 52 |
| G27459U | 52 |
| U27555A | 52 |
| C27765U | 52 |
| C28045U | 52 |
| U28114C | 52 |
| C28153U | 52 |
| U28176C | 52 |
| U28251G | 52 |
| G28378A | 52 |
| G28396A | 52 |
| G28806U | 52 |
| C29218U | 52 |
| U29584C | 52 |
| C29640U | 52 |
| C2061U | 51 |
| C2268U | 51 |
| C2842U | 51 |
| C3619U | 51 |
| A4870G | 51 |
| G5539A | 51 |
| C5643U | 51 |
| C8937U | 51 |
| C8982U | 51 |
| C10078U | 51 |
| C12534U | 51 |
| G15380U | 51 |
| C17339U | 51 |
| C17439U | 51 |
| C18312U | 51 |
| G21123U | 51 |
| C23854U | 51 |
| C25317U | 51 |
| C26936U | 51 |
| U27539C | 51 |
| G27664A | 51 |
| U28094C | 51 |
| C28507U | 51 |
| G29227U | 51 |
| C29580U | 51 |
| G347A | 50 |
| C560U | 50 |
| C1150U | 50 |
| C1427U | 50 |
| C1722U | 50 |
| A1846G | 50 |
| C2704U | 50 |
| G3728U | 50 |
| U4640C | 50 |
| G5008U | 50 |
| C6354U | 50 |
| A8837G | 50 |
| C9996U | 50 |
| C13517U | 50 |
| C13554U | 50 |
| A14004U | 50 |
| A16534G | 50 |

| Nucleotide mutation | Total number of genomes harboring the nucleotide mutation |
|---|---|
| G17280U | 50 |
| C17678U | 50 |
| U19584C | 50 |
| C21054U | 50 |
| U22334C | 50 |

**Supplementary Table S3. List of the amino acid mutations by decreasing order of frequency among the 61,397 genomes**

| Amino acid mutations | Total number of genomes harboring the nucleotide mutation causing the amino acid change |
|---|---|
| S:D614G | 61214 |
| ORF1b:P314L | 61098 |
| ORF1a:T3255I | 43710 |
| S:T478K | 40333 |
| N:R203K | 28423 |
| N:G204R | 28113 |
| ORF1a:S3675- | 27902 |
| ORF1a:G3676- | 27902 |
| S:P681H | 27345 |
| S:S477N | 26072 |
| S:L452R | 25591 |
| S:N501Y | 25345 |
| S:H655Y | 23653 |
| S:D796Y | 23545 |
| N:R32- | 23525 |
| ORF9b:E27- | 23525 |
| ORF9b:N28- | 23525 |
| N:E31- | 23522 |
| E:T9I | 23505 |
| S:N679K | 23502 |
| S:N969K | 23502 |
| N:P13L | 23491 |
| ORF1a:P3395H | 23490 |
| S:Q954H | 23475 |
| M:A63T | 23373 |
| ORF9b:P10S | 23307 |
| S:G339D | 23079 |
| ORF9b:A29- | 23049 |
| N:S33- | 23041 |
| ORF1b:I1566V | 22782 |
| S:S375F | 22423 |
| S:S373P | 22367 |
| S:N764K | 22206 |
| N:D377Y | 21564 |
| M:I82T | 21437 |
| S:P681R | 21399 |
| ORF1b:G662S | 21334 |
| ORF3a:S26L | 21310 |
| S:Q498R | 21278 |
| ORF8:D119- | 21229 |
| S:T19R | 21223 |
| ORF8:F120- | 21208 |
| N:R203M | 21186 |
| S:Y505H | 21181 |
| N:D63G | 21097 |
| ORF9b:T60A | 21092 |
| ORF1b:P1000L | 20932 |
| S:E484A | 20829 |
| S:R158G | 20669 |
| S:F157- | 20667 |
| S:E156- | 20666 |
| S:D950N | 20439 |
| ORF1a:V2930L | 20174 |
| ORF1a:T3646A | 20160 |
| ORF1a:A1306S | 20138 |
| ORF1a:P2046L | 20127 |
| ORF7b:T40I | 20110 |
| ORF1b:A1918V | 20107 |
| ORF7a:T120I | 20030 |
| ORF1a:F3677- | 19854 |
| ORF7a:V82A | 19827 |
| N:G215C | 19563 |
| S:G142D | 19558 |
| ORF1a:P2287S | 19493 |

| Amino acid mutations | Total number of genomes harboring the nucleotide mutation causing the amino acid change |
|---|---|
| S:H69- | 18454 |
| S:V70- | 18453 |
| M:Q19E | 18041 |
| S:K417N | 17690 |
| ORF3a:T223I | 15615 |
| S:A27S | 15607 |
| ORF1b:R1315C | 15476 |
| S:T19I | 15443 |
| S:P25- | 15425 |
| S:P26- | 15425 |
| S:L24- | 15420 |
| ORF1a:G1307S | 15406 |
| S:V213G | 15375 |
| ORF1a:T842I | 15370 |
| ORF1a:S135R | 15357 |
| ORF1a:T3090I | 15292 |
| S:R408S | 15228 |
| S:T376A | 15145 |
| S:D405N | 15108 |
| ORF1b:T2163I | 15086 |
| S:S371F | 15083 |
| S:Q493R | 14983 |
| N:S413R | 14612 |
| S:T95I | 13111 |
| ORF1a:L3027F | 12920 |
| S:Y144- | 11965 |
| ORF3a:Q57H | 9525 |
| ORF6:D61L | 9522 |
| ORF1b:L829I | 9029 |
| N:Q9L | 8991 |
| ORF9b:S6C | 8987 |
| ORF1a:L3201F | 8684 |
| S:A67V | 8208 |
| ORF1a:A2710T | 8079 |
| ORF1a:I3758V | 8062 |
| S:L981F | 8057 |
| ORF1a:L3674- | 8047 |
| S:N856K | 8044 |
| S:V143- | 7973 |
| S:G142- | 7941 |
| S:T547K | 7941 |
| ORF1a:K856R | 7934 |
| S:Y145D | 7914 |
| M:D3G | 7670 |
| ORF1a:S2083- | 7540 |
| ORF1a:L2084I | 7538 |
| S:S371L | 7107 |
| S:G496S | 6748 |
| ORF1a:M3087I | 6502 |
| ORF1b:V767L | 6502 |
| ORF1b:A176S | 6501 |
| S:N440K | 6501 |
| N:A376T | 6322 |
| ORF1b:E1184D | 6299 |
| S:N211- | 6228 |
| ORF1b:K1141R | 6171 |
| S:L212I | 6154 |
| N:M234I | 6081 |
| S:F486V | 5875 |
| M:D3N | 5351 |
| S:R346K | 4003 |
| ORF8:Q27* | 3930 |
| ORF8:R52I | 3930 |
| ORF1a:T1001I | 3901 |

| Amino acid mutations | Total number of genomes harboring the nucleotide mutation causing the amino acid change |
|---|---|
| S:T716I | 3897 |
| N:D3L | 3890 |
| ORF1a:A1708D | 3878 |
| S:A570D | 3870 |
| S:D1118H | 3870 |
| S:S982A | 3867 |
| N:S235F | 3860 |
| ORF8:Y73C | 3831 |
| ORF1a:I2230T | 3759 |
| S:214:EPE | 3191 |
| ORF8:A65S | 3041 |
| ORF1a:E87D | 2554 |
| ORF1a:E1724D | 2527 |
| S:P251L | 2460 |
| ORF9b:D16G | 2358 |
| S:A222V | 2162 |
| ORF1a:K261N | 1966 |
| S:G446S | 1732 |
| ORF1a:A2529V | 1729 |
| ORF1a:V1887I | 1564 |
| ORF1b:Q866R | 1562 |
| ORF3a:D155Y | 1488 |
| ORF1a:M85- | 1399 |
| ORF1a:V84- | 1389 |
| ORF1a:T265I | 1334 |
| ORF3a:H78Y | 1321 |
| ORF8:K68* | 1304 |
| ORF1a:A3209V | 1283 |
| ORF1a:T1822I | 1274 |
| S:L5F | 1256 |
| ORF1a:V665I | 1181 |
| ORF1a:T3750I | 1180 |
| ORF1a:P1786L | 1176 |
| ORF1a:P1640L | 1174 |
| ORF1a:Q2702H | 1170 |
| ORF1a:V3718A | 1127 |
| ORF1a:A498V | 1125 |
| ORF1a:A3220V | 1097 |
| ORF1b:E1341K | 1076 |
| S:Q677H | 1059 |
| ORF8:E64* | 998 |
| ORF1a:H83- | 983 |
| ORF7a:R118G | 943 |
| ORF1a:A2909S | 939 |
| ORF1a:K141- | 934 |
| ORF1a:S142- | 934 |
| ORF1a:F143- | 934 |
| ORF1a:G82- | 925 |
| ORF1a:L3606F | 922 |
| ORF3a:A33S | 884 |
| S:T719I | 862 |
| ORF1b:T1050N | 836 |
| N:T205I | 778 |
| ORF1b:Q813H | 773 |
| S:A701V | 771 |
| N:A220V | 770 |
| ORF7a:P45L | 739 |
| ORF3a:D27Y | 712 |
| ORF1b:T1540I | 699 |
| ORF1b:R188Q | 662 |
| ORF1a:K3353R | 644 |
| ORF1a:M3862I | 644 |
| ORF1a:V86- | 628 |
| ORF3a:W131C | 608 |

| Amino acid mutations | Total number of genomes harboring the nucleotide mutation causing the amino acid change |
|---|---|
| ORF1b:M1596I | 592 |
| ORF1b:R1813H | 587 |
| N:Q418H | 571 |
| S:E484K | 560 |
| ORF1a:Q556K | 559 |
| N:G214- | 556 |
| ORF1b:P1000Q | 553 |
| S:L18F | 550 |
| ORF3a:L101P | 545 |
| ORF1b:K1383R | 532 |
| ORF3a:S171L | 532 |
| S:W64R | 522 |
| N:P151S | 519 |
| S:S98F | 515 |
| ORF1a:K1655N | 509 |
| ORF9b:V30L | 499 |
| N:G30- | 486 |
| N:S327L | 476 |
| ORF9b:M26- | 476 |
| S:D80Y | 474 |
| ORF9b:A29I | 467 |
| N:S33F | 463 |
| ORF3a:N144K | 453 |
| ORF1a:C3059F | 447 |
| ORF1b:P218L | 444 |
| N:S186Y | 444 |
| S:G181V | 441 |
| ORF8:I121- | 435 |
| N:E136D | 429 |
| ORF1b:M1156I | 428 |
| ORF7a:E41D | 427 |
| ORF1a:L3829F | 422 |
| ORF1a:P80S | 421 |
| ORF7b:L11F | 414 |
| S:R346T | 411 |
| ORF1a:S944L | 407 |
| ORF1a:P309L | 400 |
| ORF3a:103:P | 397 |
| ORF1b:M1197I | 393 |
| ORF1a:Y2281H | 378 |
| ORF3a:G224C | 377 |
| ORF1a:S538P | 376 |
| ORF1a:Q2034H | 376 |
| ORF1b:K2557R | 376 |
| ORF1a:M85V | 368 |
| ORF1a:T1597I | 363 |
| ORF1b:F685Y | 362 |
| ORF7a:F101- | 359 |
| S:V1264L | 355 |
| ORF3a:D27H | 353 |
| ORF6:R20S | 353 |
| ORF7a:R80I | 353 |
| ORF8:V5I | 351 |
| S:Y145H | 348 |
| ORF1a:T3461A | 347 |
| S:V3G | 345 |
| ORF1b:S2198I | 340 |
| ORF1a:V3475F | 337 |
| ORF1a:T1567I | 336 |
| S:L242- | 335 |
| ORF7a:I100- | 335 |
| S:A243- | 334 |
| E:P71L | 331 |
| N:H300Y | 330 |

| Amino acid mutations | Total number of genomes harboring the nucleotide mutation causing the amino acid change |
|---|---|
| ORF1a:Q3777R | 328 |
| ORF1a:T945I | 327 |
| ORF1a:D1167N | 327 |
| S:L241- | 326 |
| ORF1a:Q3346K | 324 |
| S:D80A | 323 |
| S:D215G | 323 |
| S:A1078S | 322 |
| ORF1a:L2084F | 321 |
| ORF1a:A2994V | 316 |
| ORF1b:P255T | 315 |
| ORF7a:P99- | 311 |
| ORF1a:P1803S | 310 |
| ORF1a:G1125C | 309 |
| S:D1084Y | 309 |
| ORF7a:Y97- | 308 |
| ORF7a:S98- | 307 |
| ORF1a:V3595A | 305 |
| S:V213A | 304 |
| S:D215N | 304 |
| S:T240I | 303 |
| S:210:IV | 302 |
| S:N211R | 301 |
| S:Q613H | 301 |
| ORF1a:T4164A | 300 |
| S:L212G | 300 |
| N:G204P | 300 |
| ORF3a:P42L | 299 |
| ORF1a:N1662S | 298 |
| N:D377V | 292 |
| ORF1a:A2123V | 290 |
| ORF1b:E1264D | 289 |
| S:R1091H | 288 |
| ORF1b:P85L | 287 |
| ORF1b:T1637I | 286 |
| ORF3a:Q213K | 285 |
| ORF1a:I114T | 283 |
| ORF7a:L96- | 283 |
| ORF1a:A2909V | 282 |
| ORF1a:A3023T | 281 |
| ORF7a:E95- | 281 |
| ORF1a:G519S | 280 |
| ORF7a:Q94- | 280 |
| ORF3a:L140F | 279 |
| ORF1a:T1605I | 278 |
| M:A2S | 276 |
| ORF8:A14T | 275 |
| S:V1104L | 274 |
| ORF8:G77C | 273 |
| N:V270L | 268 |
| ORF1a:D1167Y | 267 |
| ORF3a:L41F | 260 |
| S:F643L | 258 |
| ORF1a:A138V | 257 |
| ORF1b:S1273L | 257 |
| ORF3a:V202L | 257 |
| ORF1a:H1113Y | 255 |
| S:S704L | 249 |
| ORF8:C83G | 248 |
| ORF1a:V2629I | 245 |
| ORF1a:A2554T | 244 |
| ORF1a:K1247N | 243 |
| S:T250I | 243 |
| ORF1a:A3623S | 242 |

| Amino acid mutations | Total number of genomes harboring the nucleotide mutation causing the amino acid change |
|---|---|
| ORF1a:V3718F | 241 |
| ORF1a:T2648I | 239 |
| S:A522S | 236 |
| ORF1b:R164C | 235 |
| S:C1250F | 232 |
| S:T29A | 230 |
| ORF1b:T2548I | 229 |
| ORF7a:L116F | 227 |
| ORF3a:L53F | 226 |
| ORF1a:T4175I | 225 |
| ORF1b:G1129V | 224 |
| ORF1a:P2018S | 219 |
| ORF3a:W69C | 218 |
| ORF3a:T151I | 217 |
| N:S194L | 216 |
| M:H125Y | 215 |
| ORF3a:G100C | 210 |
| ORF1b:V2120L | 209 |
| ORF3a:L106F | 205 |
| ORF1b:D152Y | 204 |
| ORF7a:T14I | 204 |
| S:L1049I | 203 |
| ORF3a:S165F | 203 |
| ORF1a:D2026G | 202 |
| S:T299I | 202 |
| ORF8:H17Y | 202 |
| ORF1b:A941V | 201 |
| ORF9b:P10F | 200 |
| ORF1a:P1921L | 198 |
| ORF8:F86L | 195 |
| M:R158L | 192 |
| N:S187L | 192 |
| ORF1a:Q3966R | 191 |
| S:T859I | 191 |
| S:K1191N | 191 |
| ORF3a:P240L | 191 |
| ORF8:V62L | 191 |
| ORF3a:R134C | 188 |
| N:V72I | 188 |
| ORF1a:T2264I | 187 |
| S:A701S | 187 |
| ORF1b:H1087Y | 186 |
| S:R190S | 186 |
| ORF1a:T1567A | 184 |
| ORF1b:A2575V | 184 |
| ORF1a:E1363G | 182 |
| ORF1b:Y1247C | 182 |
| ORF1a:M2259I | 181 |
| ORF1b:D815Y | 181 |
| ORF1a:T1543I | 180 |
| S:A1020S | 179 |
| ORF3a:V77F | 179 |
| S:L452Q | 177 |
| ORF1a:Q1009R | 176 |
| ORF7a:H47Y | 176 |
| ORF1a:V1747I | 174 |
| S:K417T | 172 |
| ORF7a:Q62* | 172 |
| ORF9b:R32L | 171 |
| S:D138Y | 170 |
| ORF1a:M2380I | 169 |
| ORF8:I121L | 169 |
| ORF1a:D1547Y | 168 |
| ORF1a:T2007I | 168 |

| Amino acid mutations | Total number of genomes harboring the nucleotide mutation causing the amino acid change |
|---|---|
| ORF1a:A3456V | 168 |
| S:V1176F | 168 |
| ORF1a:Q563K | 167 |
| ORF3a:Q38R | 167 |
| N:T391I | 167 |
| S:M153I | 166 |
| ORF8:F120L | 166 |
| ORF1a:K1795Q | 165 |
| ORF1b:G2297S | 165 |
| ORF1b:D2541Y | 165 |
| ORF9b:S50L | 165 |
| N:S416L | 164 |
| ORF1b:R1315L | 163 |
| N:S413I | 163 |
| ORF1a:G400C | 162 |
| ORF8:V62M | 162 |
| N:L230F | 162 |
| ORF1a:A260S | 160 |
| ORF1a:H2092Y | 160 |
| ORF1b:V2178F | 160 |
| ORF3a:S74F | 160 |
| N:P199L | 160 |
| ORF1a:P1158L | 159 |
| ORF7b:C41F | 159 |
| ORF1a:N2596S | 156 |
| ORF1a:L730F | 155 |
| N:S193I | 155 |
| ORF3a:G172R | 154 |
| ORF1a:R24C | 153 |
| ORF1a:A1649T | 153 |
| N:R195I | 153 |
| ORF8:A65V | 152 |
| ORF8:L95F | 152 |
| ORF1b:P1570L | 151 |
| ORF1b:T2432I | 151 |
| ORF8:C102F | 151 |
| ORF1a:S302F | 150 |
| S:N658S | 150 |
| ORF1a:K1895N | 149 |
| S:K529R | 149 |
| S:I68T | 148 |
| ORF3a:A31T | 147 |
| ORF3a:H204L | 147 |
| ORF3a:Q116H | 146 |
| ORF1a:L454F | 144 |
| ORF1a:K1202N | 143 |
| S:Q173H | 142 |
| S:F888L | 142 |
| S:L1265F | 141 |
| ORF1a:T1881I | 140 |
| ORF7b:F13- | 140 |
| ORF1a:V306F | 139 |
| ORF1a:T2069I | 138 |
| ORF3a:S195P | 138 |
| S:T1027I | 137 |
| ORF1a:A591V | 135 |
| ORF1a:T814I | 135 |
| ORF1b:H1213Y | 135 |
| S:L176F | 135 |
| S:N460K | 135 |
| ORF1b:L2523F | 133 |
| S:P26S | 133 |
| S:I1018V | 133 |
| ORF1b:V345L | 132 |

| Amino acid mutations | Total number of genomes harboring the nucleotide mutation causing the amino acid change |
|---|---|
| ORF3a:L15F | 132 |
| ORF3a:G49V | 131 |
| ORF1a:S1188L | 129 |
| ORF8:S67F | 129 |
| N:D128Y | 129 |
| ORF1a:A1283V | 128 |
| ORF1a:A3571V | 128 |
| ORF8:G66- | 128 |
| ORF1b:S220G | 127 |
| ORF7a:Q76L | 127 |
| ORF1b:P314F | 126 |
| E:S55F | 126 |
| ORF1b:T730I | 125 |
| N:D343G | 125 |
| ORF1a:A2129V | 124 |
| ORF1a:T4217I | 124 |
| ORF7a:Q62- | 124 |
| ORF9b:P51L | 124 |
| ORF1a:N460Y | 123 |
| ORF1a:A1352V | 123 |
| ORF1a:P1640S | 123 |
| ORF8:E92K | 123 |
| ORF3a:V112F | 122 |
| ORF1a:T4174I | 121 |
| ORF1b:Y264H | 121 |
| N:D3Y | 121 |
| N:S202N | 121 |
| ORF1a:K120N | 120 |
| ORF9b:T60I | 120 |
| S:G181R | 119 |
| ORF1a:K3839R | 118 |
| ORF1a:S4398L | 117 |
| ORF3a:S166L | 117 |
| ORF6:D6G | 116 |
| ORF7a:P84S | 116 |
| ORF1a:T882I | 115 |
| ORF1b:V426I | 115 |
| ORF1a:C1114F | 114 |
| ORF1a:F2780L | 114 |
| ORF1b:N1191S | 114 |
| ORF9b:T83I | 114 |
| ORF1a:L2039F | 113 |
| ORF1a:A2320V | 113 |
| ORF3a:S253P | 113 |
| ORF6:F2- | 113 |
| ORF1a:S2500F | 112 |
| S:G75V | 112 |
| ORF1a:P971L | 111 |
| ORF1a:P3952S | 111 |
| ORF7a:T61I | 111 |
| ORF3a:T89I | 110 |
| ORF3a:N257- | 110 |
| ORF8:K68E | 110 |
| N:P80R | 110 |
| ORF9b:Q77E | 110 |
| ORF1b:P909L | 109 |
| S:T20N | 109 |
| ORF7a:G38* | 109 |
| ORF1b:A2132V | 108 |
| ORF1a:D589Y | 107 |
| ORF1a:A656V | 107 |
| ORF1a:V1550I | 107 |
| ORF1a:L1559F | 107 |
| ORF1a:K1720N | 107 |

| Amino acid mutations | Total number of genomes harboring the nucleotide mutation causing the amino acid change |
|---|---|
| ORF1a:S2083N | 107 |
| ORF1b:L1681F | 107 |
| ORF3a:V48F | 107 |
| N:M317T | 107 |
| ORF1a:S2535L | 106 |
| ORF1a:L3715F | 106 |
| ORF1b:P2321S | 106 |
| S:D215Y | 106 |
| S:D1260N | 106 |
| N:S2- | 106 |
| N:P368T | 106 |
| ORF1a:G645S | 105 |
| ORF1a:G3617C | 105 |
| N:A12G | 105 |
| ORF9b:H9D | 105 |
| ORF1a:I3341V | 104 |
| ORF1b:V1271L | 104 |
| S:I850L | 104 |
| ORF1a:M1312I | 103 |
| ORF3a:K16T | 103 |
| ORF3a:A103V | 103 |
| ORF9b:I5T | 103 |
| ORF1a:L1853F | 102 |
| E:V62F | 102 |
| ORF1b:H1550Y | 101 |
| ORF1b:M1581I | 101 |
| ORF9b:K40R | 101 |
| ORF1b:V1407F | 100 |
| ORF1b:G2180V | 100 |
| S:S939F | 100 |
| ORF8:L84S | 99 |
| ORF1a:N4051S | 98 |
| ORF1b:P1975S | 98 |
| ORF1b:L2213F | 98 |
| ORF3a:V259L | 98 |
| ORF7b:V21I | 98 |
| N:T135I | 98 |
| N:T379I | 98 |
| ORF1a:L1955F | 97 |
| ORF1a:T2495I | 97 |
| ORF1a:V4216I | 97 |
| ORF1b:S1182L | 97 |
| ORF1b:M1573I | 97 |
| ORF1b:R2613N | 97 |
| ORF1b:V2685F | 97 |
| ORF1a:V649F | 96 |
| ORF1a:M3752I | 96 |
| ORF1b:I214V | 96 |
| ORF1b:M620I | 96 |
| S:Q52R | 96 |
| S:G1219V | 96 |
| ORF1a:K247N | 95 |
| ORF1a:M321I | 95 |
| ORF1a:H1160Y | 95 |
| ORF1b:A86V | 95 |
| ORF1b:P1967S | 95 |
| S:S255F | 95 |
| ORF1a:A339V | 94 |
| ORF1a:T2283I | 94 |
| ORF1a:T2300I | 94 |
| ORF1b:A302S | 94 |
| S:S943P | 94 |
| S:A1087S | 94 |
| ORF7a:V104F | 94 |

| Amino acid mutations | Total number of genomes harboring the nucleotide mutation causing the amino acid change |
|---|---|
| ORF1a:T2106I | 93 |
| ORF1b:A2028S | 93 |
| S:D88H | 93 |
| S:S254F | 93 |
| S:S494P | 93 |
| ORF3a:G18D | 93 |
| ORF3a:A54S | 93 |
| ORF1a:L642F | 92 |
| ORF1b:S2689G | 92 |
| S:W152L | 92 |
| N:P365S | 92 |
| ORF1a:P2046S | 91 |
| S:K150N | 91 |
| ORF3a:V228A | 91 |
| N:K370R | 91 |
| ORF1a:T1496I | 90 |
| S:T76I | 90 |
| E:L21F | 90 |
| N:A208S | 90 |
| ORF1a:N278S | 89 |
| ORF1a:K564N | 89 |
| ORF1a:T1854I | 89 |
| ORF1a:P1862L | 89 |
| ORF1a:A3143V | 89 |
| ORF1b:V248F | 89 |
| S:F157L | 89 |
| S:T859N | 89 |
| ORF8:S67- | 89 |
| ORF1a:T727I | 88 |
| ORF1a:L3915F | 88 |
| ORF9b:Q34L | 88 |
| ORF1a:P1921S | 87 |
| ORF1a:S2224F | 87 |
| N:A152S | 87 |
| ORF1a:L451F | 86 |
| ORF1a:V779F | 86 |
| ORF1a:G1073V | 86 |
| ORF1b:L2017F | 86 |
| S:S12F | 86 |
| ORF3a:A99S | 86 |
| ORF7a:L102- | 86 |
| ORF8:Q18* | 86 |
| N:K373N | 86 |
| ORF1a:G1119S | 85 |
| ORF1a:D2136G | 85 |
| ORF1a:Y2141- | 85 |
| ORF1a:V2642I | 85 |
| ORF1a:D3196G | 85 |
| ORF1a:L3201I | 85 |
| ORF1a:T4065I | 85 |
| ORF1b:A2268V | 85 |
| S:T883I | 85 |
| S:T1117I | 85 |
| ORF7a:A105V | 85 |
| ORF1a:P62S | 84 |
| ORF1a:V337I | 84 |
| ORF1a:A516V | 84 |
| ORF1a:P4197L | 84 |
| ORF7a:T28I | 84 |
| ORF1a:A690V | 83 |
| ORF1a:T2016I | 83 |
| ORF1a:A3615V | 83 |
| ORF1b:T1774I | 83 |
| ORF1b:P2531S | 83 |

| Amino acid mutations | Total number of genomes harboring the nucleotide mutation causing the amino acid change |
|---|---|
| S:P1162L | 83 |
| ORF3a:Y215H | 83 |
| ORF3a:V256- | 83 |
| ORF9b:P10L | 83 |
| ORF1a:S391F | 82 |
| ORF1a:T4161I | 82 |
| ORF3a:Y107H | 82 |
| ORF7a:F63- | 82 |
| ORF7a:E121* | 82 |
| ORF1a:V2613F | 81 |
| N:T362I | 81 |
| ORF1a:P1213L | 80 |
| S:D1260Y | 80 |
| ORF1a:S245Y | 79 |
| ORF1a:L1130F | 79 |
| ORF1a:I3052V | 79 |
| ORF1a:P3371S | 79 |
| ORF1a:T4083M | 79 |
| ORF1b:T1511I | 79 |
| ORF7a:S60- | 79 |
| ORF7a:T61- | 79 |
| N:T24I | 79 |
| N:W330L | 79 |
| ORF1a:G1100S | 78 |
| ORF1a:A1204V | 78 |
| ORF1a:A2554S | 78 |
| ORF1b:D2179A | 78 |
| ORF1b:T2537I | 78 |
| ORF3a:S40L | 78 |
| N:P151L | 78 |
| N:Q289H | 78 |
| ORF1a:S558F | 77 |
| ORF1a:T1246I | 77 |
| S:212:SGR | 77 |
| ORF7a:A105S | 77 |
| ORF1a:R124C | 76 |
| ORF1a:A429V | 76 |
| ORF1a:I2501T | 76 |
| ORF1b:D2067G | 76 |
| S:H49Y | 76 |
| S:L841F | 76 |
| N:T393I | 76 |
| ORF1a:T1754I | 75 |
| ORF1a:N3540S | 75 |
| ORF1b:V344I | 75 |
| ORF1b:M2414I | 75 |
| S:A262S | 75 |
| ORF3a:T175I | 75 |
| ORF7a:F59- | 75 |
| N:M210I | 75 |
| ORF1a:P804S | 74 |
| ORF1b:A1521S | 74 |
| ORF1b:A1643V | 74 |
| S:P1162S | 74 |
| ORF3a:G49C | 74 |
| ORF3a:A99V | 74 |
| ORF7a:L17F | 74 |
| ORF1a:A1204T | 73 |
| ORF1a:A1997V | 73 |
| S:R78M | 73 |
| S:D936Y | 73 |
| ORF3a:G18V | 73 |
| ORF3a:A110S | 73 |
| ORF8:E106* | 73 |

| Amino acid mutations | Total number of genomes harboring the nucleotide mutation causing the amino acid change |
|---|---|
| ORF1a:L681F | 72 |
| ORF1a:T1788M | 72 |
| ORF1a:I2873V | 72 |
| ORF1a:Y3204H | 72 |
| ORF1a:S3384L | 72 |
| ORF1b:R2613C | 72 |
| S:G446V | 72 |
| S:L452M | 72 |
| ORF3a:K21N | 72 |
| ORF3a:V88L | 72 |
| ORF3a:V255- | 72 |
| N:A90S | 72 |
| ORF9b:E86D | 72 |
| ORF1a:Y947- | 71 |
| ORF1a:E948- | 71 |
| ORF1a:K2078E | 71 |
| ORF1b:A927V | 71 |
| ORF1b:R1052K | 71 |
| S:T22I | 71 |
| ORF9b:M8I | 71 |
| ORF1a:A1043V | 70 |
| ORF1a:P1054L | 70 |
| ORF1a:T1682I | 70 |
| ORF1a:T2152I | 70 |
| ORF1a:I3853V | 70 |
| ORF1b:T861I | 70 |
| ORF1b:P1427S | 70 |
| ORF1b:Q1546H | 70 |
| ORF3a:A23S | 70 |
| N:G238C | 70 |
| ORF9b:T24I | 70 |
| ORF1a:P286L | 69 |
| ORF1a:H1500Y | 69 |
| ORF1a:T1754N | 69 |
| ORF1a:S2193F | 69 |
| ORF1a:A2584V | 69 |
| ORF1b:V2073L | 69 |
| ORF1b:A2143V | 69 |
| ORF1b:Q2247H | 69 |
| ORF8:W45L | 69 |
| ORF1a:A903V | 68 |
| ORF1a:P971S | 68 |
| ORF1a:I1551T | 68 |
| ORF1a:L3199S | 68 |
| ORF1a:A3392V | 68 |
| ORF1b:P970L | 68 |
| ORF1b:D1572N | 68 |
| S:S151- | 68 |
| S:S221L | 68 |
| S:D253G | 68 |
| ORF3a:A110V | 68 |
| ORF7a:I103- | 68 |
| ORF1a:P380L | 67 |
| ORF1a:S2103F | 67 |
| ORF1a:S2242F | 67 |
| ORF1a:T3488I | 67 |
| ORF1a:I4049V | 67 |
| ORF1b:P1664S | 67 |
| S:W152- | 67 |
| S:V705I | 67 |
| ORF1a:T1168I | 66 |
| ORF1a:A2097V | 66 |
| ORF1b:P1570S | 66 |
| ORF1b:A2222T | 66 |

| Amino acid mutations | Total number of genomes harboring the nucleotide mutation causing the amino acid change |
|---|---|
| ORF1b:A2431V | 66 |
| ORF1b:T2488M | 66 |
| S:N439K | 66 |
| ORF1a:T403I | 65 |
| ORF1a:S443F | 65 |
| ORF1a:T1035I | 65 |
| ORF1a:T2093I | 65 |
| ORF1a:V3689M | 65 |
| ORF1b:D1183Y | 65 |
| S:T250N | 65 |
| S:T547I | 65 |
| S:D614N | 65 |
| ORF7a:I88S | 65 |
| ORF1a:E767G | 64 |
| ORF1a:D996G | 64 |
| ORF1a:T2274I | 64 |
| ORF1b:T76I | 64 |
| S:L938F | 64 |
| ORF7a:V24F | 64 |
| ORF7a:T39I | 64 |
| ORF8:H17Q | 64 |
| N:Q83R | 64 |
| ORF9b:K80E | 64 |
| ORF1a:V108I | 63 |
| ORF1a:T1000I | 63 |
| ORF1a:S2900L | 63 |
| ORF1b:I97V | 63 |
| ORF3a:D155H | 63 |
| ORF3a:V244I | 63 |
| ORF9b:A57V | 63 |
| ORF1a:G3278S | 62 |
| ORF1b:V311L | 62 |
| ORF1b:G1561S | 62 |
| ORF1b:I2687T | 62 |
| S:A684V | 62 |
| S:V1264A | 62 |
| ORF3a:T24I | 62 |
| ORF3a:T221K | 62 |
| ORF1a:L2146F | 61 |
| ORF1a:L3338F | 61 |
| ORF1a:A3610V | 61 |
| ORF1b:E735D | 61 |
| ORF1b:E1288D | 61 |
| ORF1b:L1504F | 61 |
| ORF1b:M2667I | 61 |
| S:W258L | 61 |
| S:T732I | 61 |
| ORF3a:G254V | 61 |
| N:P142S | 61 |
| N:P168S | 61 |
| N:R209I | 61 |
| ORF9b:T72I | 61 |
| ORF1a:H110Y | 60 |
| ORF1a:M1378I | 60 |
| ORF1a:Q1813P | 60 |
| ORF1b:H2388Y | 60 |
| ORF3a:G172C | 60 |
| ORF7a:P34L | 60 |
| ORF7a:P34S | 60 |
| ORF8:S54L | 60 |
| N:R203I | 60 |
| N:Q289L | 60 |
| ORF1a:Q314E | 59 |
| ORF1a:T1022I | 59 |

| Amino acid mutations | Total number of genomes harboring the nucleotide mutation causing the amino acid change |
|---|---|
| ORF1a:A3956T | 59 |
| ORF1b:G1093S | 59 |
| ORF1b:T1336I | 59 |
| ORF1b:L1351F | 59 |
| ORF1b:P1427L | 59 |
| ORF1b:T2040I | 59 |
| ORF3a:A72S | 59 |
| ORF7b:L14- | 59 |
| N:P364L | 59 |
| ORF1a:H417Y | 58 |
| ORF1a:T999I | 58 |
| ORF1a:A1397V | 58 |
| ORF1a:L2105F | 58 |
| ORF1a:A2637T | 58 |
| ORF1b:Q348H | 58 |
| ORF1b:A1219S | 58 |
| ORF1b:T1522I | 58 |
| N:M322I | 58 |
| ORF1a:V1056L | 57 |
| ORF1a:M1586I | 57 |
| ORF1a:T3284I | 57 |
| ORF1a:T4087I | 57 |
| ORF1a:M4241I | 57 |
| ORF1b:Y1464C | 57 |
| S:P1263L | 57 |
| ORF3a:L108F | 57 |
| ORF1a:S34T | 56 |
| ORF1a:I841V | 56 |
| ORF1a:Q1140R | 56 |
| ORF1a:P1692S | 56 |
| ORF1a:S2285F | 56 |
| ORF1a:K2589R | 56 |
| ORF1a:L4345F | 56 |
| ORF1a:A4357V | 56 |
| ORF3a:W131R | 56 |
| ORF7a:V71L | 56 |
| ORF7a:T115I | 56 |
| ORF8:F120V | 56 |
| N:T366I | 56 |
| ORF1a:A1234S | 55 |
| ORF1a:H2799Y | 55 |
| ORF1b:M115I | 55 |
| ORF1b:T284I | 55 |
| ORF1b:M1397I | 55 |
| ORF1b:A1521V | 55 |
| ORF1b:P2313S | 55 |
| S:H245Y | 55 |
| S:E484Q | 55 |
| ORF7b:E3* | 55 |
| ORF8:A55V | 55 |
| ORF1a:S212L | 54 |
| ORF1a:M3189I | 54 |
| ORF1b:V711I | 54 |
| ORF1b:A1895V | 54 |
| ORF1b:D2090Y | 54 |
| ORF1b:G2610D | 54 |
| ORF7a:V104- | 54 |
| ORF8:A65- | 54 |
| ORF8:Y79C | 54 |
| ORF9b:P3L | 54 |
| ORF1a:E640D | 53 |
| ORF1a:S2024L | 53 |
| ORF1a:P2605S | 53 |
| ORF1a:T2967I | 53 |

| Amino acid mutations | Total number of genomes harboring the nucleotide mutation causing the amino acid change |
|---|---|
| ORF1a:P4337S | 53 |
| ORF1b:V1538L | 53 |
| ORF1b:P1727L | 53 |
| ORF1b:F2314L | 53 |
| S:P272L | 53 |
| ORF3a:A23V | 53 |
| ORF3a:A54V | 53 |
| ORF1a:R398C | 52 |
| ORF1a:I567V | 52 |
| ORF1a:H2659Y | 52 |
| ORF1b:L1220F | 52 |
| ORF1b:M1693T | 52 |
| ORF1b:A2585S | 52 |
| S:R21I | 52 |
| S:A520S | 52 |
| ORF7a:E22D | 52 |
| ORF7a:F54L | 52 |
| ORF8:I74T | 52 |
| ORF8:T87I | 52 |
| N:G178V | 52 |
| N:Q229H | 52 |
| ORF9b:R32H | 52 |
| ORF9b:G38D | 52 |
| ORF1a:A599V | 51 |
| ORF1a:A668V | 51 |
| ORF1a:A1793V | 51 |
| ORF1a:A2891V | 51 |
| ORF1a:T2906I | 51 |
| ORF1b:S638I | 51 |
| ORF1b:A1291V | 51 |
| S:F490S | 51 |
| S:S1252F | 51 |
| ORF7a:E91K | 51 |
| ORF7b:L4F | 51 |
| ORF9b:A75V | 51 |
| ORF1a:V28I | 50 |
| ORF1a:R99C | 50 |
| ORF1a:H388Y | 50 |
| ORF1a:A486V | 50 |
| ORF1a:G1155C | 50 |
| ORF1a:S2030L | 50 |
| ORF1a:I2858V | 50 |
| ORF1a:S3244L | 50 |
| ORF1b:T17I | 50 |
| ORF1b:K179N | 50 |
| ORF1b:S1023G | 50 |
| ORF7b:E39* | 50 |
| ORF8:A51V | 50 |
| N:A211T | 50 |
| N:T417I | 50 |
